# Potent and selective repression of *SCN9A* by engineered zinc finger repressors for the treatment of neuropathic pain

**DOI:** 10.1101/2024.09.06.609976

**Authors:** Mohammad Samie, Toufan Parman, Mihika Jalan, Jisoo Lee, Patrick Dunn, Jason Eshleman, Dianna Baldwin Vidales, Josh Holter, Brian Jones, Yonghua Pan, Marina Falaleeva, Sarah Hinkley, Alicia Goodwin, Tammy Chen, Sumita Bhardwaj, Alex Ward, Michael Trias, Anthony Chikere, Madhura Som, Yanmei Lu, Sandeep Yadav, Kathleen Meyer, Bryan Zeitler, Jason Fontenot, Amy Pooler

## Abstract

Peripheral neuropathies are estimated to affect several million patients in the US with no long-lasting therapy currently available. In humans, the Nav1.7 sodium channel, encoded by the *SCN9A* gene, is involved in a spectrum of inherited neuropathies, and has emerged as a promising target for analgesic drug development. The development of a selective Nav1.7 inhibitor has been challenging, in part due to structural similarities among other Nav channels. Here we present preclinical studies for the first genomic medicine approach using engineered zinc finger repressors (ZFRs) specifically targeting human/nonhuman primate (NHP) *SCN9A*. AAV-mediated delivery of ZFRs in human iPSC-derived neurons resulted in 90% reduction of *SCN9A* with no detectable off-target activity. To establish proof-of-concept, a ZFR targeting the mouse *Scn9a* was assessed in the SNI neuropathic pain mouse model, which resulted in up to 70% repression of *Scn9a* in mouse DRGs and was associated with reduction in pain hypersensitivity as measured by increased mechanical- and cold-induced pain thresholds. AAV-mediated intrathecal delivery of ZFR in NHPs demonstrated up to 60% repression of *SCN9A* in bulk DRG tissue and on single-cell levels in the nociceptors. The treatment was well tolerated in NHPs, and no dose-limiting findings were observed four weeks after a single intrathecal injection. Taken together, our results demonstrate that AAV-delivered ZFR targeting the *SCN9A* gene is promising and supports further development as a potential therapy for peripheral neuropathies.

## INTRODUCTION

Peripheral neuropathy, a common chronic pain condition, is recognized as one of the most difficult pain syndromes to manage(*1, 2*). It results from direct damage to somatosensory neurons,(*3*) and it may manifest in numerous ways including small fiber neuropathy, diabetic peripheral neuropathy, trigeminal neuralgia, and post herpetic neuralgia. These conditions, all of which are characterized by severe damage to peripheral neurons, are estimated to affect several million patients in the USA(*4*). Importantly, patients with peripheral neuropathies are usually refractory to common pain medications. Given the high unmet need and lack of durable treatment options, there is an urgent need to develop novel and long-lasting therapeutics for treatment of chronic peripheral neuropathies.

Nerve injury activates a population of primary afferent sensory neurons called nociceptors, which are responsible for translating noxious stimuli into electrical signals. These signals, in the form of action potentials, are then transmitted from the periphery to the cortex, where the brain can perceive the signal as pain (*5*). Voltage-gated sodium channels (Nav channels) are responsible for initiating and propagating the action potential to the brain and are synthesized in the soma of nociceptors (*6*). The soma resides within the dorsal root ganglia (DRG) located close to the intervertebral foramina at the cervical, thoracic, lumbar, and sacral spinal cord levels. Nav channels are major molecular regulators of neuronal excitability and are widely expressed in the central and peripheral nervous systems as well as within cardiac and skeletal muscles. To date, nine different Nav channels (Nav1.1-Nav1.9) have been identified (*7, 8*). Of the nine known Nav channels in humans, Nav1.7, Nav1.8, and Nav1.9 have been found to be principally expressed in nociceptors and are known to be associated with pain (*6*). Among these, Nav1.7 is involved in a spectrum of inherited human pain disorders (*9*). Gain-of-function mutations in the *SCN9A* gene that encodes for Nav1.7 have been linked to pain in inherited erythromelalgia (IEM) (*10, 11*) and paroxysmal extreme pain disorder (PEPD), with which patients exhibit a severe excessive pain phenotype (*12*). Alternatively, loss-of-function mutations in *SCN9A* are associated with a clinical condition called congenital insensitivity to pain (CIP), in which individuals exhibit complete loss of pain sensation (*13*). Further studies have shown that Nav1.7 defines the threshold of the action potential and amplifies small depolarizing inputs, acting as the major molecular regulator of signal transduction in nociceptive neurons (*14, 15*), thus having an essential and non-redundant role in transduction and transmission of pain signaling following noxious stimuli (*16*). Patients with CIP do not display any motor, cognitive, sympathetic, or gastrointestinal deficiencies, and except for the loss of pain sensation and in some cases anosmia, they generally exhibit normal sensory functions (*13, 17*). This strongly suggests that Nav1.7 is predominantly involved in pain sensation and establishes Nav1.7 as a promising target candidate for analgesic drug development. Although preclinical (*18–21*) and clinical (*22*) studies have shown that repression of Nav1.7 leads to reduction of pain, the development of a selective Nav1.7 small molecule inhibitor has been challenging. This is primarily due to similarities in protein sequence and structure among the various Nav channels, resulting in unwanted side effects and failure in clinical trials(*23–26*).

To avoid the complexities associated with targeting the Nav1.7 protein, we aimed to regulate Nav1.7 more specifically by epigenetically repressing the endogenous expression of *SCN9A* using engineered ZFRs. Zinc fingers (ZFs) are naturally occurring transcription factor proteins that have primarily evolved to regulate eukaryotic gene expression epigenetically and represent the most abundant and diverse class of DNA binding proteins in the human genome (*27*). Over the past several years, ZFRs were used to repress the expression of different genes involved in various neurological disorders in different preclinical models including chronic pain(*20*), Huntington’s disease (*28*) and Alzheimer’s disease (*29*). ZFRs hold several advantages over other genomic medicine approaches; they do not nick or induce double strand breaks in endogenous genomic DNA and their components are of human origin, minimizing immunogenicity risk in patients. Furthermore, consistent with their dominant role as part of about 350 different native human Krüppel-associated box (KRAB) proteins, ZFRs often outperform bacterially sourced CRISPR formats in epigenetic transcriptional modulation of human genes (*20, 30*) and because they are native to human body, ZFRs are expect to be generally well tolerated in patients. Lastly, ZFR compact size allows for unrestricted packaging into adeno associated virus (AAV) vectors, thereby enabling the generation of efficient, specific, and long-lasting down-regulation of *SCN9A* expression through a single administration.

Here we present a novel genomic medicine approach for targeting conserved sequences within *SCN9A* in human and NHP genomes. To establish proof-of-concept, efficacy of ZFRs was assessed in a mouse model of neuropathic pain. Mouse surrogate ZFR (mZFR) targeting the mouse *Scn9a* gene were designed and evaluated in the spared nerve injury (SNI) mouse model of neuropathic pain. AAV9-mediated mZFR delivery resulted in up to 70% bulk repression of *Scn9a* in mouse DRG tissue. Repression of *Scn9a* was associated with a significant reduction in pain 1 month after a single IT-L administration as measured by the mechanical- and cold-induced pain threshold in the SNI model of neuropathic pain. To develop the clinical candidate, we designed and screened a library containing several hundred ZFRs targeting the human/NHP *SCN9A* gene in human neuronal cell lines. The lead ZFR was then packaged into an AAV9 vector and tested in human iPSC-derived GABAergic and iPSC-derived sensory neurons *in vitro*. ZFRs were selected that repressed the expression of *SCN9A* by more than 90% over a wide dose range with no detectable off-target activity as measured by global transcriptomic analysis, including selective repression of Nav1.7 mRNA. We evaluated a candidate human ZFR in a 1-month pharmacology and safety study in cynomolgus monkeys, assessing ZFR expression and concomitant *SCN9A* repression in the dorsal root ganglion (DRG), showing repression of *SCN9A* at 40-60% in all DRG levels. A significant reduction of *SCN9A* mRNA was seen in NHP nociceptor neurons by using single-cell analysis. The lead ZFR was well tolerated at all doses tested with no ZFR-related dose-limiting findings up to 1 month after administration of a single bolus intrathecal-lumbar (IT-L) dose to cynomolgus monkeys, supporting the progression of the program into GLP IND-enabling toxicology studies. Taken together, these results demonstrate that an AAV-delivered ZFR targeting the Nav 1.7 gene is promising and supports further development as a potential therapy for peripheral neuropathy indications in patients.

## RESULTS

### NHP/Human *SCN9A* targeting ZFRs potently and selectively reduces human *SCN9A* levels in iPSC-derived neurons without altering the expression of other Nav channels *in vitro*

Considering the central role of Nav1.7 in transmitting pain signals to the brain, we hypothesized that reducing Nav1.7 via selective repression of the *SCN9A* gene using ZFRs would reduce the expression of Nav1.7 in neurons, and thereby reduce neuropathic pain in peripheral neuropathies. ZFRs are engineered by combining a designed zinc finger array with a KRAB domain (*31*). The zinc finger array mediates site-specific binding to the *SCN9A* gene, and the KRAB domain represses the endogenous expression of *SCN9A* transcript, leading to a reduction in Nav1.7 protein (Fig. 1A).

**Figure 1.**
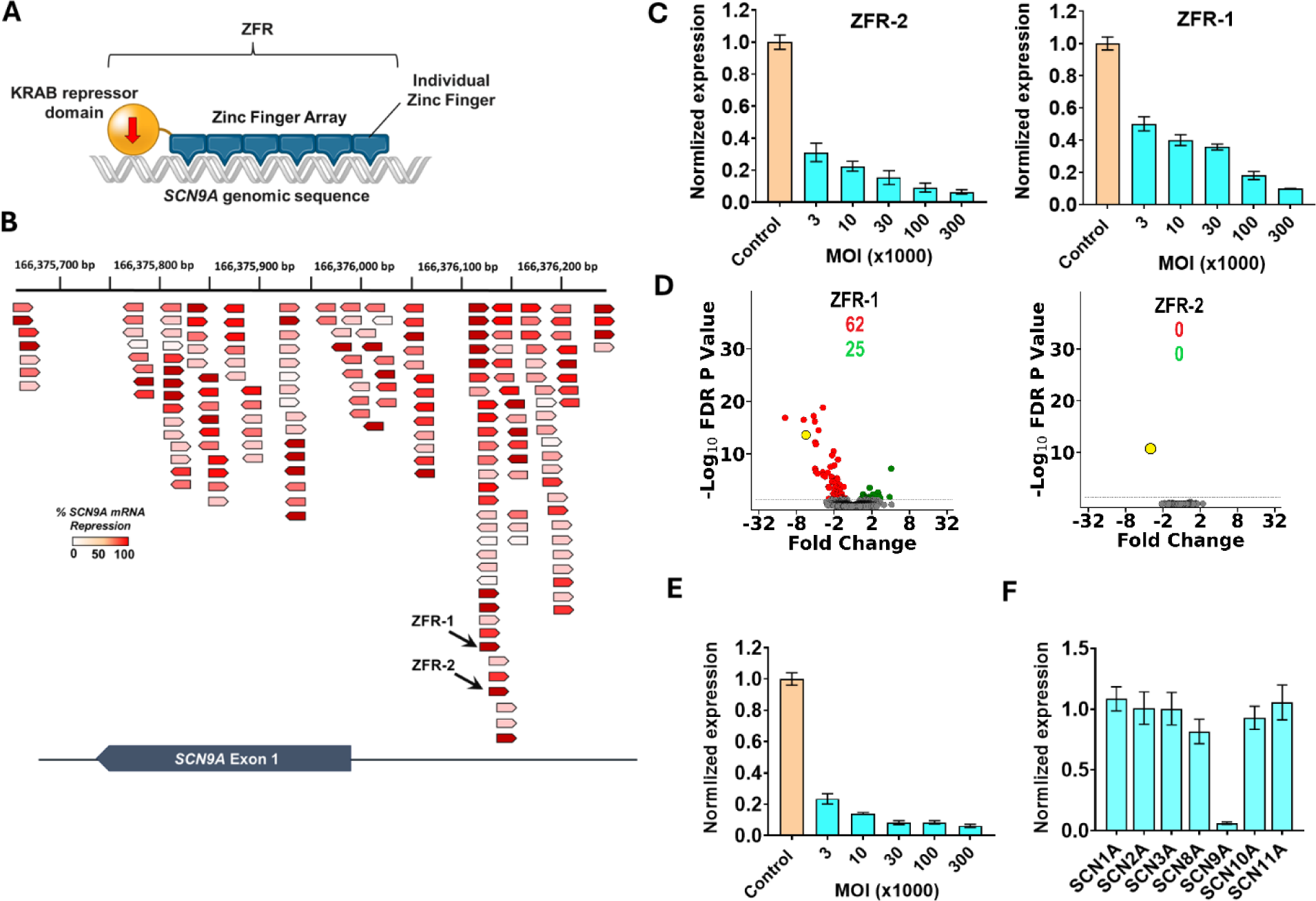
*SCN9A* targeting hZFR specifically reduces human *SCN9A* levels in iPSC derived neurons without influencing the expression of other Nav channels *in vitro*. **(A**) Engineered ZFRs consist of two main domains; a designed zinc finger array domain which is specifically binding to the *SCN9A* genomic sequence, and a KRAB transcriptional repressor domain from the human KOX1 protein. The ZFR selectively binds to near the *SCN9A* transcription start site, recruits the KAP1 epigenetic modification complex via the KRAB domain, and represses the expression of *SCN9A*. (**B**) Schematic of binding locations around the human *SCN9A* TSS and *SCN9A* mRNA repression levels of 200 ZFRs tested in SK-N-MC cells. Each arrow indicates a ZFR and its orientation (sense or antisense) regarding *SCN9A* exon 1. The heat map (red-white) illustrates *SCN9A* expression levels following ZFR treatment. The arrow color red indicates over 90% *SCN9A* repression, while white represents no repression of *SCN9A*. The numbers on top represent the genomic location on chromosome 12. (**C)** Significant reduction of *SCN9A* transcriptional levels in a dose-dependent manner in iPSC-derived GABAergic neurons following AAV6 mediated transduction of ZFR-1 or ZFR-2. Doses (MO1) 3E3, 1E4, 3E4, 1E5, and 3E5. Mean ± SD. (**D)** mRNA microarray assessment (volcano plots) of iPSC-derived GABAergic neurons 30 days post-transduction with ZFR-1 or ZFR-2 packed into AAV6 and delivered to cells at 1E5 MOI. Each dot represents the mean fold change compared to control-treated cells for a given gene (x value) and the associated P value (y value). Gene expression profiles are calculated based on FDR adjusted P value >0.05. Red dots represent genes that are down regulated, green dots represent genes that are upregulated, and yellow dot indicates *SCN9A* gene. 6 biological replicates. (**E)** Significant reduction of *SCN9A* transcriptional levels in a dose-dependent manner in iPSC-derived sensory neurons following AAV6 mediated transduction of ZFR-2 (hZFR). Doses (MO1) 3E3, 1E4, 3E4, 1E5, and 3E5. Mean ± SD. (**F)** hZFR only represses *SCN9A* and no other Nav channels. iPSC-derived sensory neurons were transduced with AAV6-hZFR at 3E5 MOI and cells were recovered 7 days post transduction and processed for gene analysis. ZFRs specifically repressed *SNC9A* and minimally affected the expression of other Nav channels at high dose, Mean ± SD.

To identify potent and specific ZFRs, we assembled a library containing several hundred ZFRs targeting conserved sequences between humans and NHPs located on the *SCN9A* transcription start site (TSS) near *SCN9A* exon 1 on chromosome 12 (Fig. 1B). The screening of ZFRs was performed in the SK-N-MC human neuroepithelial cell line using ZFR mRNA nucleofection. SK-N-MC cells express *SCN9A* transcripts at high levels and are thus appropriate for testing ZFRs that reduce *SCN9A* expression. Some ZFRs failed to decrease *SCN9A* transcript levels, whereas others exhibited a potent repression of *SNC9A* transcript with more than 90% repression (Fig. 1B). Based on repression profiles, several ZFRs were selected for further evaluation in iPSC-derived GABAergic neurons. AAV6 has exhibited the greatest ability to transduce a wide range of primary cells in culture compared to other serotypes (*32*), thus AAV6 was used for the *in vitro* characterization of ZFRs in neurons. AAV6-mediated delivery of ZFR-1 and ZFR-2 yielded a significant and dose-dependent repression of *SCN9A* mRNA levels in iPSC-derived GABAergic neurons (Fig. 1C), with minimal effect on the expression of the housekeeping gene A*TP5B* (fig. S1), demonstrating that the ZFRs can potently reduce *SCN9A* transcript levels in iPSC-derived neurons *in vitro*.

To evaluate the potential ZFR off-target impact on global gene expression, microarray analysis was conducted on total RNA isolated from human iPSC-derived GABAergic neurons transduced with ZFRs. Three weeks after transduction with 1E5 multiplicity of infection (MOI) of the ZFRs packed into AAV6 vectors, cells were harvested and analyzed using human Clariom S microarrays. The Clariom S microarray contains probes targeting more than 21,000 genes, allowing the conduct of a comprehensive analysis of global gene expression in neurons. In addition to *SCN9A*, ZFR-1 differentially regulated the expression of 80 other genes, while ZFR-2 specifically repressed the expression of *SCN9A* mRNA and did not regulate the expression of any other genes in the iPSC-derived neurons (Fig. 1D). ZFR-2 (hereinafter referred to as hZFR) was further characterized in iPSC-derived sensory neurons (which are the target neuronal population for neuropathic pain treatment). AAV6-mediated transduction of hZFR in sensory neurons resulted in a potent and dose-dependent repression of *SCN9A* 10 days after transduction (Fig. 1E). Furthermore, the expression of other Nav channels in the iPSC-derived sensory neurons was evaluated following transduction with high hZFR doses using RT-qPCR probes designed specifically for each Nav channel. hZFR potently repressed *SCN9A* with no significant effect on expression of other Nav channels in iPSC-derived sensory neurons (Fig. 1F). Altogether, data obtained from iPSC-derived neurons indicates that hZFR potently and selectively targets *SCN9A in vitro*.

### Potent repression of *Scn9a* in the spared nerve injury (SNI) mouse model of neuropathic pain

As the target genome sequences encoding the Nav1.7 sodium channel differ between mouse *Scn9A* and NHP/human *SCN9A*, surrogate mZFRs were generated to demonstrate the effectiveness of ZFRs *in vivo* in a mouse neuropathic pain model. mZFRs were screened in mouse Neuro2A (N2A) neural crest-derived cell line and a lead mZFR with potent repression (fig. S2A) was selected for further off-target evaluation in mouse primary cortical neurons (MCNs). One week after transducing the MCNs with AAV6-mZFR with 3E4 MOI, the total RNA isolated from the MCNs was analyzed using mouse Clariom S microarray. The lead mZFR exhibited no off-target activity. However, *Scn9a* is not differentially regulated in MCNs because it is not actively expressed in these central nervous system neurons; thus no changes in the *Scn9a* levels were observed (fig. S2B). Prior to conducting pain assessment studies, the on-target activity of AAV9 mediated delivery of mZFR (AAV9-mZFR) was evaluated in lumbar DRGs of C57BL/6 mice 4 weeks after a single IT-L administration (Fig. 2A). mZFR was under the control of a neuronal specific human Synapsin (Syn) promoter and it was highly expressed in the bulk lumbar DRG analysis 4 weeks after treatment (fig. S2C). While a significant repression of *Scn9a* was observed in bulk lumbar DRGs, no statistical upregulation of neuroinflammation marker (*Iba1*) or downregulation of neuronal marker (*Rbfox3*) compared to the vehicle group were observed, demonstrating lack of neuroinflammatory responses or neuronal loss in the lumbar DRG 1 month after AAV9-mZFR treatment in mice (Fig. 2B-D).

**Figure 2.**
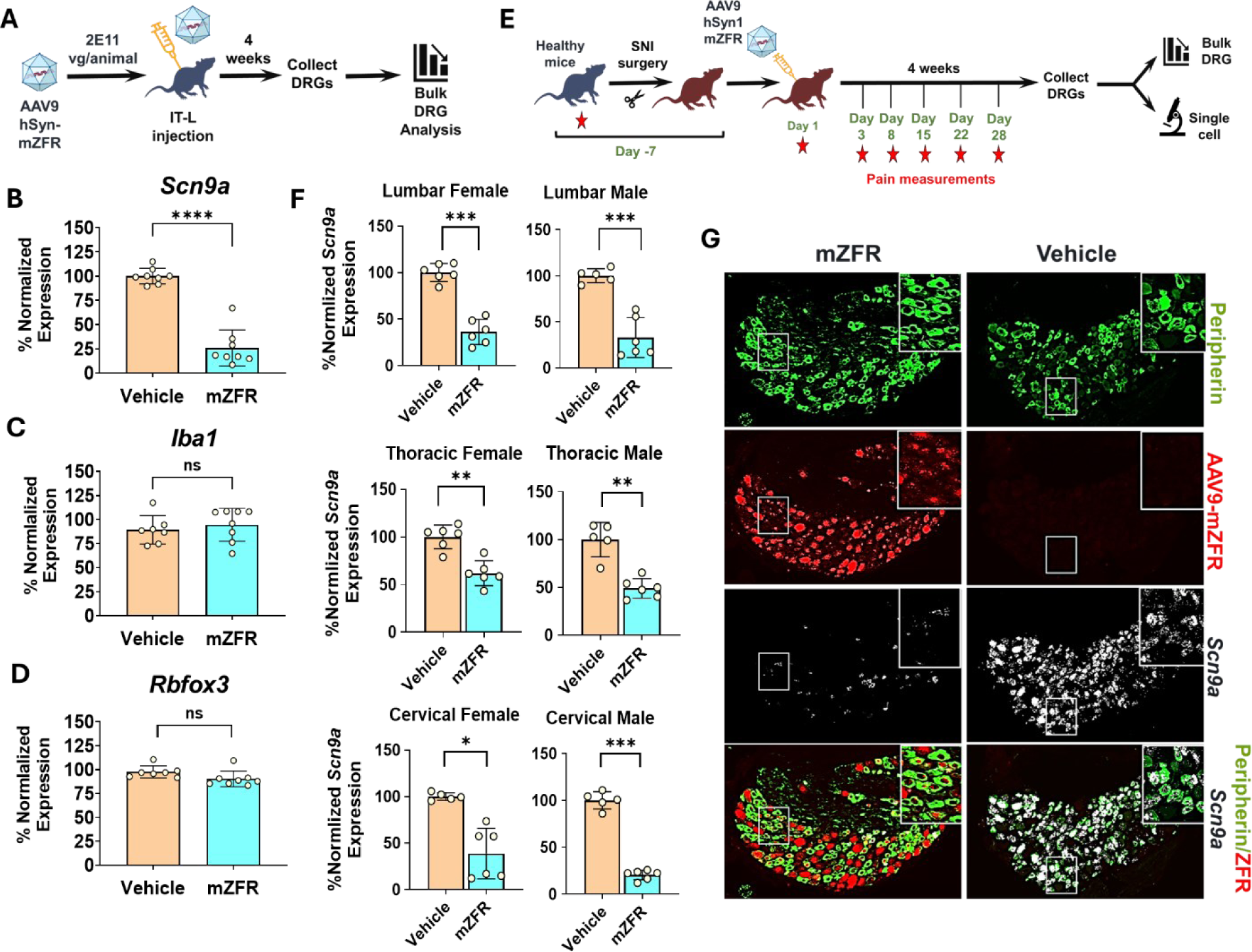
ZFR mediated repression of mouse *Scn9A* in a mouse model of neuropathic pain. **(A)** Overview and timeline of target engagement and tolerability study in WT mice using mZFR packed into AAV9. (**B)** Normalized mRNA expression of *Scn9a* in the lumbar DRG, four weeks after IT-L injection compared to the vehicle group. Mean ± SEM. (**C)** Normalized mRNA expression of *Iba1* (macroglia mediated neuroinflammatory marker), in the lumbar DRG, four weeks after IT-L injection compared to the vehicle group. Mean ± SEM. (**D)** Normalized mRNA expression of *Rbfox3* (neuronal marker) in the lumbar DRG, four weeks after IT-L injection compared to the vehicle group. Mean ± SEM. (**E)** Overview and timeline of the pain efficacy study using the SNI mouse model. Red stars indicate the time points where pain responses were measured. Mechanical and cold allodynia were measured in WT mice before they undergo SNI surgery. Seven days after the SNI surgery (Day 1), mechanical and cold allodynia were measured again, and animals were injected with AAV9-hSyn1-mZFR at 8E11 vg/animal. Pain responses were measured at different timepoints post injection (Days 3, 8, 15, 22, and 28). After four weeks, animals were sacrificed and DRGs were collected for gene expression and pathological analysis. Stars indicate the timepoints pain responses were measured. (**F)** Normalized average mRNA expression of *Scn9a* in mouse lumbar, thoracic, and cervical DRGs obtained from female and male animals. *Scn9a* levels were significantly reduced by 60-70% at the bulk lumbar and cervical DRG tissue level four weeks post IT-L injection with AAV9-mZFR. Mean ± SEM, ordinary one-way ANOVA, *\*\* P < 0.01 *** P < 0.001* **** *P* < 0.0001 compared with vehicle. **(G)** Single-cell analysis illustrating the reduction of *Scn9a* mRNA in AAV9-mZFR expressing nociceptors. Lumbar DRG from mouse were immunolabeled for peripherin (nociceptor marker – green) and hybridized with fluorescently labeled mouse specific *Scn9a* and ZFR RNA probes. Four weeks post injection, while a very high expression of *Scn9a* (white) is observed in the animals treated with the vehicle, a significant reduction of *Scn9a* mRNA is observed in peripherin positive nociceptors in animals injected with AAV9-mZFR (red).

The effectiveness of mZFR was evaluated in the Spared Nerve Injury (SNI) mouse model of neuropathic pain (Fig 2E). Results from this model provide understanding of the relationship between ZFR expression, *Scn9a* repression and efficacy in reducing neuropathic pain. The SNI model in rodents is a partial denervation model that involves cutting the common peroneal and tibial nerves while sparing the sural nerve, leading to consistent and reproducible tactile pain hypersensitivity in the skin territory of the spared intact sural nerve (*33*) (fig. S2D). The SNI model has proven to be robust, demonstrating substantial and prolonged changes in measures of mechanical sensitivity and thermal responsiveness, reminiscent of clinically described conditions for neuropathic pain disorders (*33*). Prior to surgical procedures, a pain sensitivity baseline was measured for all animals using Von Frey filaments for the assessment of mechanical-induced pain (filaments were applied from the underside of the mesh to the surface of the mouse hind paw) and a cold plate for cold-induced pain. Seven days after the SNI surgery and after collection of a baseline responses of the mouse model to mechanical- and cold-induced pain, mice (8 males and 8 females) were assigned to their respective groups and a single IT-L dose of either vehicle or AAV9-mZFR at 8E11 vg/animal was administered into the L4-L5 intervertebral space. For the sham (control) group, animals underwent surgery, but nerves were not cut (only skin opened and sutured closed) and AAV9-mZFRs were not administered. One month following dosing, animals were euthanized, DRGs were collected and evaluated for expression levels of *Scn9a* at bulk and single-cell levels. Mechanical- and cold-induced pain were measured during the course of the study to assess efficacy of mZFR (Fig. 2E). Additionally, clinical observations, body weights, and necropsy observations were performed to assess tolerability. Bulk DRG analysis using RT-qPCR demonstrated approximately 70% reduction in *Scn9a* expression in lumbar and cervical DRG levels compared to approximately 40% repression in thoracic DRGs in both male and female animals (Fig. 2F). There were no gender differences in terms of *Scn9a* repression. RNAscope in situ hybridization (ISH) was used in combination with immunohistochemistry (IHC) to assess *Scn9a* mRNA repression on a single-cell level. The nociceptors were identified with a peripherin specific antibody (green). High levels of *Scn9a* mRNA were observed in peripherin positive cells in lumbar DRG isolated from the vehicle group. A significant reduction of *Scn9a* mRNA (white) transcript was observed in peripherin-positive cells in AAV9-mZFR treated animals (Fig. 2G & fig. S2E), illustrating that mZFRs significantly reduce the expression of *Scn9a* transcript in mouse nociceptors.

### *In vivo* repression of mouse *Scn9a* reverses pain hypersensitivity in SNI mouse model of neuropathic pain

Mechanical- and cold-induced pain were measured pre-dose before SNI surgery to establish a baseline (Fig 3A-B). Prior to dose administration (7 days post SNI surgery), mechanical- and cold-induced pain were measured again to establish baseline before treatment. All mice that underwent SNI surgery demonstrated significantly lower mechanical- and cold-induced pain thresholds compared to the sham control group. This was evident by less force or temperature necessary to the affected limb to elicit a withdrawal behavior as compared to the sham group. Animals treated with a single IT-L dose of AAV9-mZFR demonstrated a statistically significant increase in mechanical-induced pain threshold compared with their respective vehicle control groups on Day 28 (Fig 3A). On Day 28, the mechanical-induced pain threshold in AAV9-mZFR animals was comparable to that of the sham group, indicating that ZFR treatment was able to restore pain responses to normal levels. Additionally, to further validate the effectiveness of AAV9-mZFR, a small-molecule analgesic drug, gabapentin (GBP), was included as a positive control. GBP was administered intraperitoneally approximately 1 hour before testing on Day 28. The timing for GBP administration was selected based on its reported time to reach maximum plasma concentration and nociceptive effectiveness. Paw withdrawal latency in response to cold-induced pain significantly increased in AAV9-mZFR-treated animals compared with the vehicle group (Fig. 3B). On Day 28, the cold-induced pain threshold in mZFR-treated animals was comparable to sham and GBP-treated animals. AAV9-mZFR was able to rescue the pain phenotype as early as 3 days post-dose in the SNI neuropathic pain model and the pain threshold gradually increased to the same level as in the sham-treated animals by Day 28 (Extended Data Fig.3).

**Figure 3.**
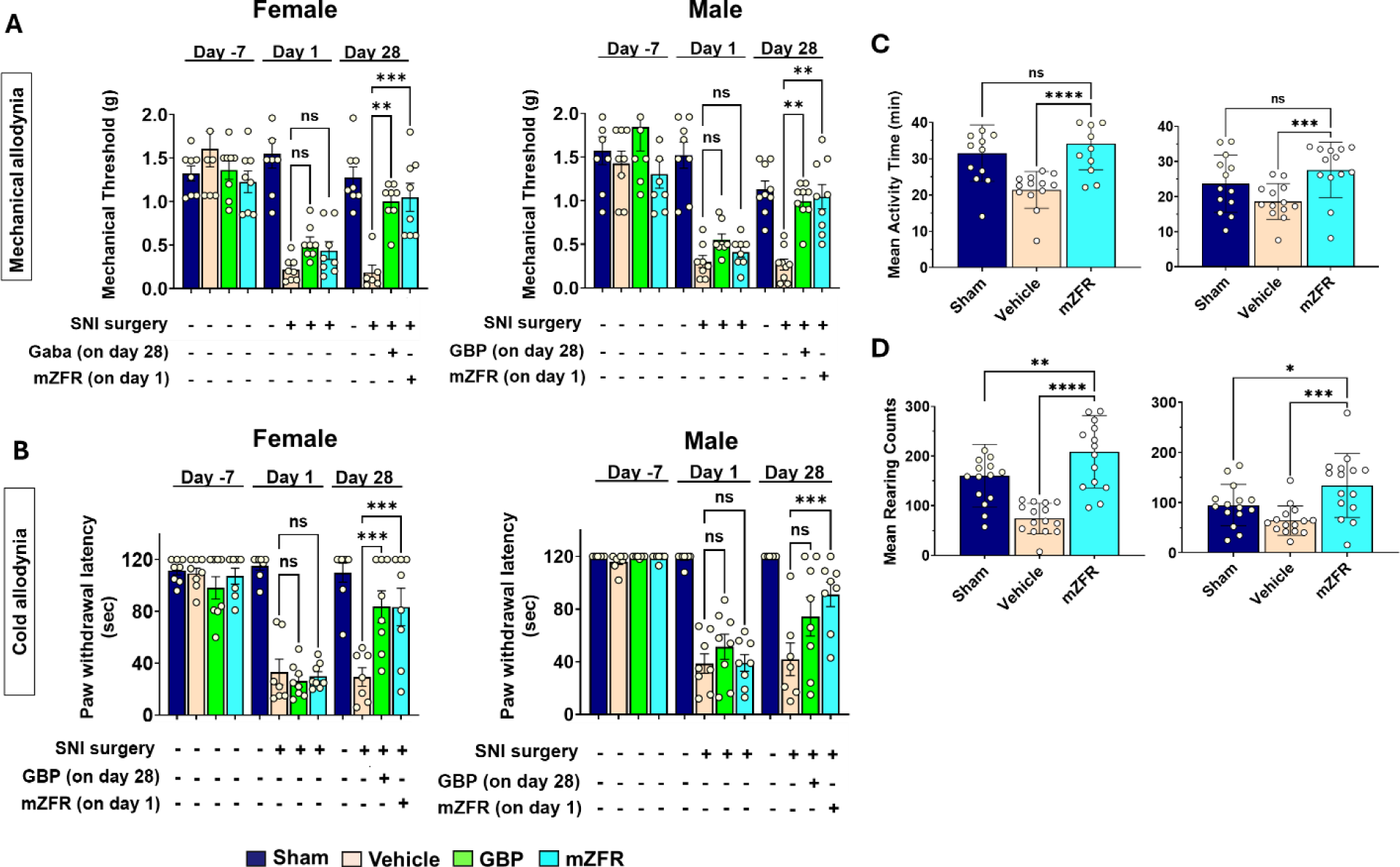
*In vivo* repression of mouse *Scn9A* reverses pain hypersensitivity in a mouse model of neuropathic pain without altering mice overall activity. **(A and B)** Mechanical (**A**) or cold (**B**) induced pain responses were measured at Day -7 (healthy mice) and then at Day 1 (7 days post SNI surgery). Twenty-eight (28) days after AAV9-mZFR IT-L injection, treated animals exhibited a significantly higher pain threshold compared to the vehicle group and at a comparable level to the sham group for both female and male animals. Gabapentin (GBP) was injected on Day 28 one hour before pain measurements. Dots represent individual animals; n = 7-8; Mean ± SEM, one-way ANOVA, ** P < 0.01 *** P < 0.001 **** P < 0.0001 compared to vehicle group. **(C & D)** SmartCage assessment was performed on SNI mice at Day 28 post IT-L injection of AAV9-mZFR. Meant activity time (**C**) and mean rearing count (**D**) were evaluated for both male and female animals. During the dark cycle, vehicle-treated animals exhibited a lower value for both parameters. Animals treated with AAV9-mZFR exhibited similar levels of activity and rearing count compared to the same group and statistically significantly higher than the vehicle group. Dots represent a time point in the dark cycle; Mean ± SEM; one-way ANOVA, ** P < 0.01 *** P < 0.001 **** P < 0.0001 compared to vehicle group.

SmartCage® analysis was utilized to evaluate any possible changes or recovery in animal movement after treatment with AAV9-mZFR. Mice are nocturnal, and their peak activity time is during the dark cycle, thus the SmartCage data were analyzed between the hours of 6 pm and 5 am. The relevant SmartCage parameters for the SNI mouse model are mean activity time (MAT) and mean rearing count (MRC). MAT measures free movement within the cage while MRC evaluates the number of times animals stand on both hind paws in a vertical upright position. Animals experiencing pain usually have less movement and thus have lower MAT and/or MRC values. In general, the vehicle group exhibited lower values for MAT and MRC compared to the sham group, which is anticipated since sham animals did not have surgical manipulation (Fig. 3C-D). MAT and MRC levels significantly increased in AAV9-mZFR treated animals compared to vehicle group and were similar to levels in sham group animals for both male and female mice (Fig. 3C-D). Altogether, data demonstrate efficacy in the SNI mouse model, showing that *in vivo* repression of mouse *Scn9a* reverses pain hypersensitivity and thereby improves movement and rearing in an SNI mouse model of neuropathic pain.

### Specific and dose dependent repression of *SCN9A* in multiple DRG levels 1 month after intrathecal-lumbar injection in NHPs

Pharmacology and target specificity of AAV9-mediated delivery of hZFR (AAV9-hZFR) was evaluated in cynomolgus monkeys in a 1-month dose range-finding (DRF) pharmacology and toxicology study following a single IT-L administration at three dose levels: 1E12, 1E13, and 9E13 vg/animal (Fig. 4A). The potency of the AAV9-hZFR was evaluated at bulk DRG levels (lumbar, thoracic, and cervical) and in nociceptors using single-cell analysis. AAV9-hZFR was expressed in a dose-dependent manner in all DRG regions analyzed (Fig. 4B), and repressed *SCN9A* expression by 40-60% at bulk DRG levels compared to the vehicle group (Fig. 4C).

**Figure 4.**
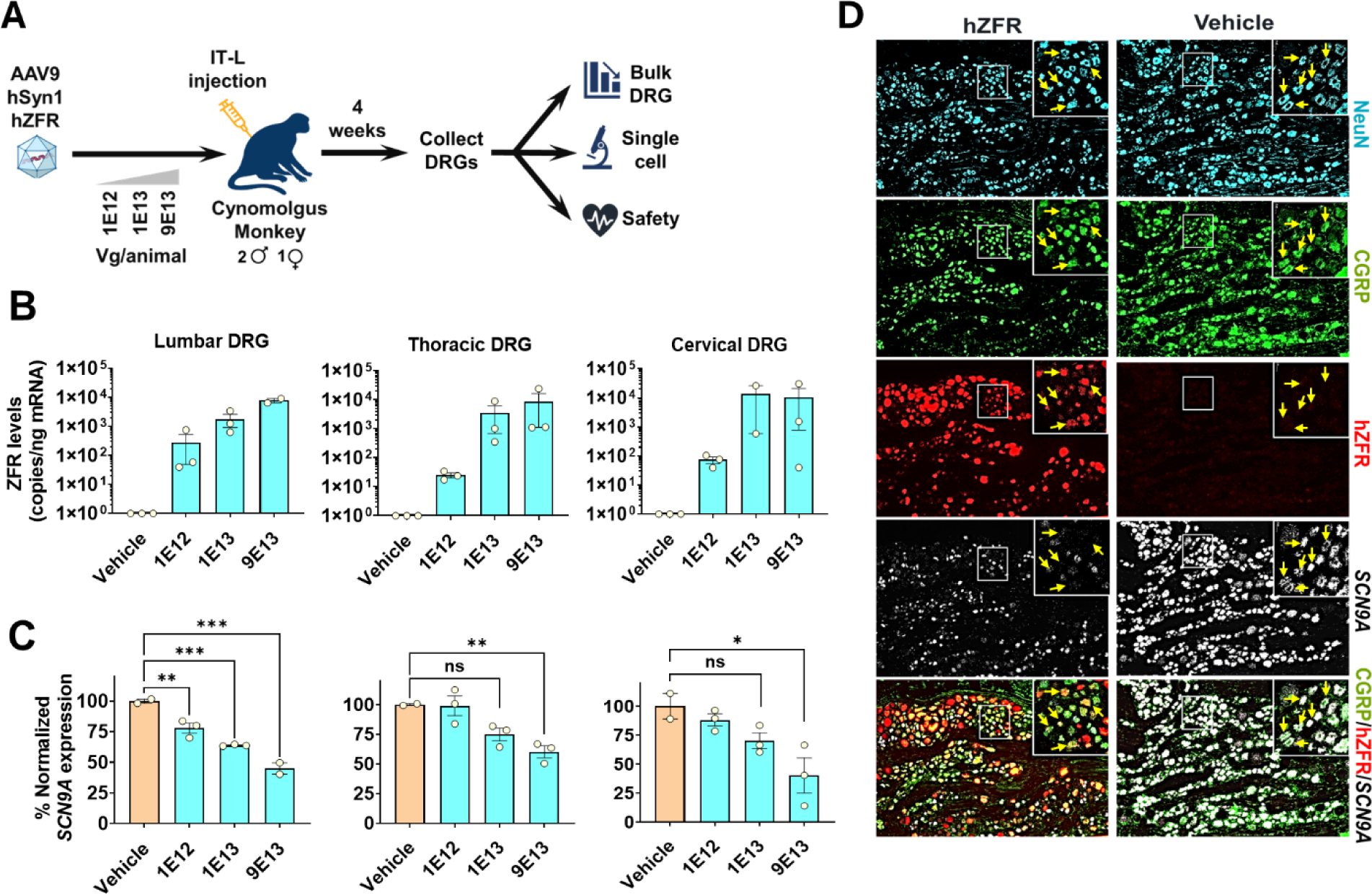
Dose dependent repression of *SCN9A* in multiple DRG levels one month after IT-L injection in NHPs. **(A)** Overview and timeline of the NHP study evaluating the potency and safety of the hZFR. **(B)** Average AAV9-hZFR expression in NHP lumbar, thoracic, and cervical DRGs, 4 weeks post IT-L injection at 3 different doses. Dots represent an individual NHP. Mean ± SEM. **(C)** Normalized average mRNA expression of *SCN9A* in NHP lumbar, thoracic, and cervical DRGs. *SCN9A* levels were reduced by 40-60% at the bulk DRG tissue level across a 100-fold dose range four weeks post injection. Dots represent an individual NHP; Mean ± SEM; one-way ANOVA, ** P < 0.01 *** P < 0.001 **** P < 0.0001 compared to vehicle group. **(D)** Single-cell analysis illustrating the reduction of *Scn9a* mRNA in AAV9-hZFR expressing nociceptors. The left column shows lumbar DRG sections obtained from the hZFR treated NHPs and right column shows sections obtained from the vehicle treated NHPs. Lumbar DRG sections from NHPs injected with the high dose (9E13 vg/animal) of AAV9-hZFR or vehicle control were immunolabeled for NeuN (neuronal marker-teal), CGRP (nociceptive marker – green) and hybridized with fluorescently labeled NHP specific *SCN9A* RNA probe (white) and ZFR RNA probe (red). Four weeks post injection, nociceptor neurons containing AAV9-hZFR (red), do not express *SCN9A* RNA compared to neurons lacking AAV9-hZFR. Yellow arrows point to NeuN/CGRP positive cells.

RNAscope ISH evaluated the ZFR-mediated repression of *SCN9A* on a single-cell level. Unlike mice, the cellular and molecular bases of nociception in NHPs are less understood and much less is known about characteristics of neuronal populations in NHP or human DRG. We tested several commercially available antibodies against proteins predicted to be expressed in nociceptors in higher species such as CGRP (CALCA), IB4 (P2X3R), and TAC1 (Substance P). Among all tested antibodies, an antibody against CGRP demonstrated consistent and clear signal, and thus selected to further characterize the expression of *SCN9A* in nociceptors in NHP lumbar DRG. It has been shown before that approximately 60% of CGRP positive neurons in human DRG are positive for *SCN9A* (*34*). An antibody against NeuN (neuronal marker) was used to assess the total neuronal population in the DRGs. These antibodies were combined with RNAscope ISH to assess *SCN9A* mRNA levels in nociceptors following hZFR injection (Fig 4D). A high level of *SCN9A* mRNA (white) was observed throughout lumbar DRGs in CGRP positive cells (green) in vehicle treated NHPs. A significant reduction of *SCN9A* mRNA transcript was observed in CGRP positive cells that were also positive for hZFR (Fig. 4D), illustrating that ZFRs can significantly reduce expression of *SCN9A* transcript in NHP DRG nociceptors. Taken together, results confirmed the pharmacologic activity of hZFR in NHPs. To confirm the specificity of hZFR *in vivo,* we evaluated the expression of other known Nav channels expressed in the DRG including Nav1.6 (*SCN8A*), Nav1.8 (*SCN10A*), and Nav1.9 (*SCN11A*), following IT-L administration of AAV9-hZFR. Supporting target specificity, no changes in expression levels of other Nav channels were observed in any DRG level at any dose, while a significant and dose-dependent repression of *SCN9A* in the bulk DRG analysis was observed 1 month post IT-L treatment with hZFR (Fig.5). Altogether, the DRF study demonstrated the potency and specificity of hZFR in NHP DRGs.

**Figure 5.**
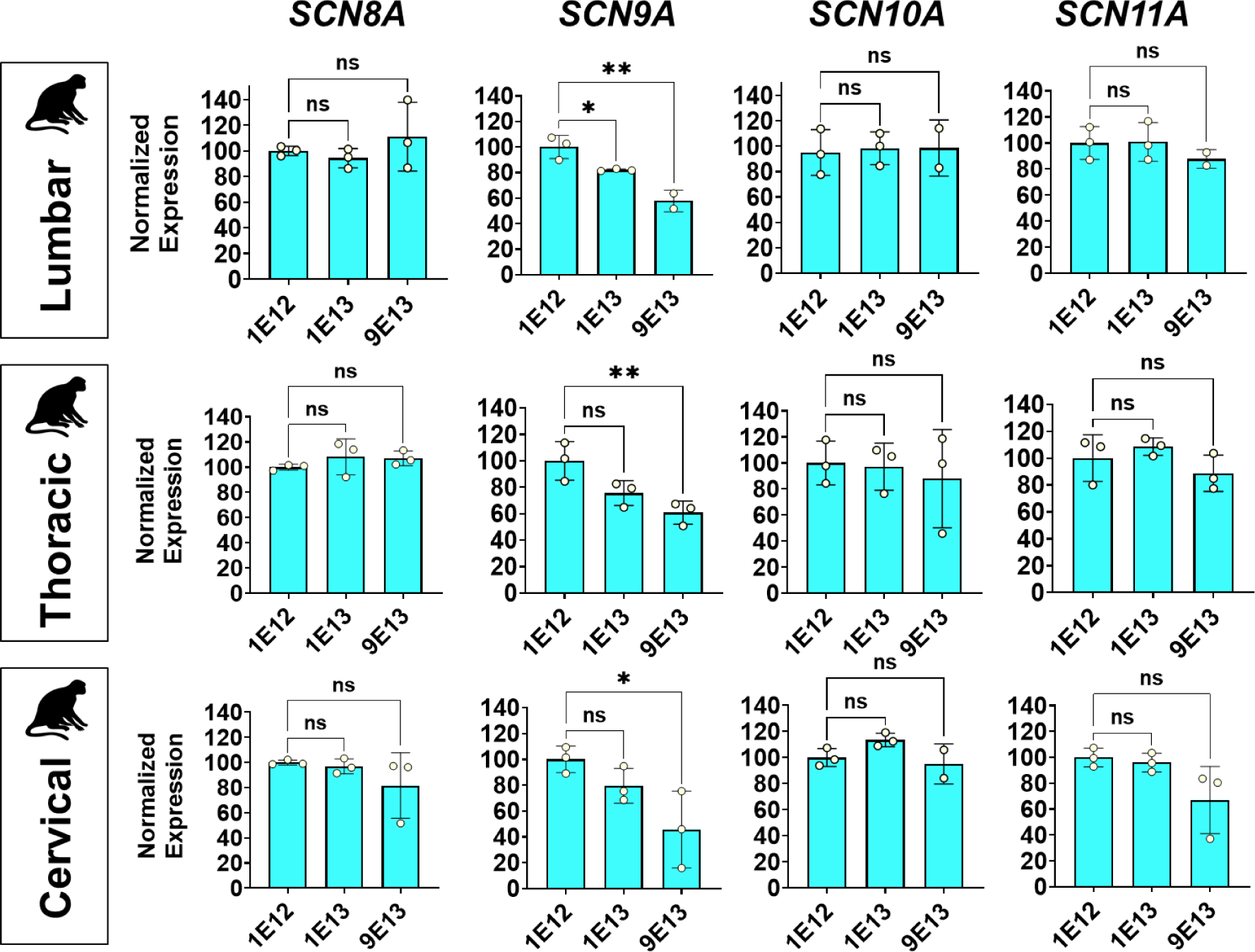
AAV9-hZFR does not influence the expression of other Nav channels in the DRGs. The percentage expression levels of *SCN8A* (Nav1.6), *SCN9A* (Nav1.7), *SCN10A* (Nav1.8), and *SCN11A* (Nav1.9) are presented for each individual DRG level that was collected and analyzed for each dose (1E12, 1E13, and 9E13 vg/animal). There was no dose-dependent repression of *SCN8A*, *SCN10A*, and *SCN11A* observed at any DRG level at any dose. Each circle represents an NHP. Mena ± SEM. one-way ANOVA, ** P < 0.01 *** P < 0.001 **** P < 0.0001 compared to vehicle group.

### AAV9 mediated delivery of hZFR demonstrated an acceptable safety profile in NHPs

hZFR was well tolerated in cynomolgus monkeys and all animals survived until scheduled necropsy. There were no AAV ZFR-related abnormal clinical signs, behavior, body weight, clinical chemistry, hematology, coagulation, urinalysis parameters or necropsy findings. Histopathology was performed for the following tissues: adrenal glands, brain, DRGs (sacral, lumbar, thoracic, and cervical), epididymis, heart, kidney, small and large intestines, liver, lung, lymph node (mandibular), ovary, pancreas, sciatic nerve, skeletal muscle, spinal cord (lumbar, thoracic, cervical), spleen, stomach, testes, thymus, trigeminal ganglia (TG), and uterus/cervix. Among all treated animals, histopathology analysis showed no treatment-related findings in the brain or the other collected tissues, except, DRGs, spinal cords, and TG and sciatic nerve.

Microscopic evaluation showed dose-related findings in the TG and DRG of minimal to mild mononuclear cell infiltration (MN) and/or minimal degeneration/necrosis of single neurons (SCN), minimal to mild axonal degeneration (AD) in the spinal cord and/or minimal MN of dorsal roots, and additionally for the high dose, minimal AD and minimal MN in the sciatic nerve. Mild severity was only noted in the high dose group for MN in DRG (lumbar, 3/3 animals; sacral, 1/3 animals), and AD in spinal cord (1/3 animal). These findings are known class-effects of AAV gene therapies in NHPs (*35–38*), and thought primarily related to over-expression of transgenes in neurons and associated effects on axons originating from these neurons(*39, 40*). Treated animals did not develop adverse neurological signs throughout the 1-month study, similar to what has been reported in the literature following AAV gene therapy administered to the CNS or systemically (*39, 40*).

## DISCUSSION

Peripheral neuropathy represents a major health burden and a globally unmet clinical need (*41*), and this type of pain is associated with greater anxiety, depression, and sleep disturbance than non-neuropathic pain (*42*). Current medications indicated for treatment of neuropathic pain may potentially provide some relief, but the effect sizes for these drugs are small, and treatments are often associated with serious adverse effects. Moreover, many patients will not obtain sufficient pain relief at tolerated doses (*1, 42–45*). Opioids are another treatment option for neuropathic pain yet can be accompanied by a range of harmful secondary side effects and addictive potential (*46*). Considering the lack of long-lasting and safe treatments for chronic neuropathic pain, there is urgent need for new drugs for chronic pain management, and genomic medicines may offer an attractive alternative approach for neuropathic pain.

Nav1.7 has gained much attention in the past couple of decades due to findings that loss or gain of function mutations in *SCN9A* lead to loss of or excess pain sensation in patients, respectively, providing genetic evidence for the Nav1.7 sodium channel as a therapeutic target for pain (*14, 15, 47*). Most attempts, however, to develop specific small molecule inhibitors targeting Nav1.7 have failed in the clinic. Several reasons have been proposed for failures, such as insufficient channel blockade (*48*) or nonoptimal clinical trial designs which led to mixed results (*22*). One of the key issues has been a lack of selectivity toward Nav1.7 and resulting adverse effects associated with inadvertently targeting other Nav channels(*23–26*). Nav channels share many sequence and structural similarities, and play critical roles in maintaining neuronal excitability in different tissues. Thus, targeting these sodium channels could possibly have unwanted clinical consequences. Nav1.1, Nav1.2, and Nav1.3 are mainly expressed in the brain and loss of function in these channels is linked to epilepsy and autism. Nav1.4 and Nav1.5 are expressed predominantly in the muscular system and are responsible for maintaining muscle cell action potentials, including in cardiac muscle cells (*8*). Nav1.6, Nav1.8, and Nav1.9 are mainly expressed in the DRG, and like Nav1.7, have been implicated in pain pathways (*49*). Recently, a selective small-molecule inhibitor targeting Nav1.8 has shown promising results in clinical trials for acute pain (*17*). However, Genome Wide Association Studies (GWAS) as well as other analyses revealed that loss of function mutations in Nav1.8 can possibly lead to Brugada syndrome (sudden death syndrome) and cardiac arrest (*50–52*), so the long-term effect and safety of Nav1.8 inhibitors in the clinic remains to be established.

Considering the wide expression and range of functions Nav channels play in the body, it is critical to develop a selective repressor for Nav1.7 for repression of neuropathic pain. Utilizing the specificity of ZFRs in targeting precise DNA sequences, we have identified unique sequences and binding sites near the human/NHP *SCN9A* TSS that are not shared with other Nav channels. Global transcriptomic analysis revealed that hZFR only represses the expression of *SCN9A,* and no other genes are differentially regulated including other Nav channels in human iPSC-derived GABAergic and sensory neurons (Fig. 1). In correlation, no changes in expression levels of Nav1.6, Nav1.8, or Nav1.9 were observed in NHP bulk DRG analysis 1 month post IT-L treatment with hZFR (Fig 5). Collectively, these data demonstrate that human ZFRs selectively and specifically target *SCN9A in vitro* and in mice and NHPs *in vivo*.

IT-L administration of hZFR to cynomolgus monkeys demonstrated ZFR mRNA expression levels capable of significantly reducing bulk *SCN9A* mRNA expression. Similar levels were seen in the SNI neuropathic pain model in mice administered mZFRs, suggesting delivery, expression and activity of ZFRs in NHPs may translate to patients and provide therapeutic benefit. Similar data were obtained at the single-cell level in NHP lumbar DRG neurons, exhibiting repression of *SCN9A* in ZFR-positive nociceptors, further demonstrating potency of ZFRs, and that low cellular levels of ZFRs are effective at reducing up to 90% of target *SCN9A* expression. The degree of inhibition of *SCN9A* needed to effectively reverse neuropathic pain in the clinic has not yet been established. Individuals, however, who are heterozygous for a loss-of-function mutation of Nav1.7 (individuals who contain only one functional allele of *SCN9A* and presumably express 50% of functional Nav1.7 protein) exhibit normal pain responses (*15*), indicating that 50% repression of Nav1.7 should be sufficient to establish a normal pain response. In the SNI neuropathic pain mouse model, we showed that 60% repression of *Scn9a* at the bulk DRG level was sufficient to fully restore normal pain responses for mechanical- and cold-induced pain. Additionally, hZFR administration induced up to 60% bulk repression of *SCN9A* in all three DRG levels 1 month after IT-L administration to NHPs, suggesting that hZFR treatment may be sufficient to reduce neuropathic pain in patients. Currently there is no reliable NHP model of neuropathic pain; thus, translatability of pain reduction from rodent to human can only be assessed during clinical trials. It should be noted that full repression of Nav1.7 function and pain responses are not desired, where complete lack of Nav1.7 expression in humans may lead to a state of insensitivity to pain where unintended self-injury, such as burning, could occur.

The proof-of-concept mouse model of neuropathic pain using mouse surrogate ZFRs was an important step in preclinical development. This model allows understanding of the relationship between ZFR mRNA expression, *Scn9a* mRNA repression and efficacy in a pain model. In cynomolgus monkeys, a single IT-L administration of AAV9-hZFR at doses up to 9E13 vg/animal resulted in similar levels of *SCN9A* repression, was well tolerated and not associated with ZFR-related adverse effects including clinical signs, body weights, clinical pathology and macroscopic observations 1 month after the treatment. Microscopic evaluation associated with AAV9-hZFR-treatment showed dose-related findings of minimal to mild histopathologic findings primarily in the DRGs as well as spinal cord, sciatic nerve and trigeminal ganglia, which has been reported for other AAV gene therapies (*35–38*). Notwithstanding the aforementioned microscopic findings, no abnormal behavior indicative of neurological dysfunction was exhibited by AAV9-hZFR-treated animals.

The DRG and spinal cord findings were consistent with AAV-delivered gene therapy and published literature for IT-L administration (*53*). Meta-analysis of 33 studies with more than 200 NHPs based on published data revealed that DRG toxicity is believed to be a class effect of AAV gene therapy, where several factors contribute to the incidence and severity of DRG histopathology findings in NHPs, including: direct administration into the CSF via intrathecal or intracisternal magna injection, and administration of dose levels greater than 1E13 vg/animal (*53*). Incidence and severity are also influenced by route of administration, vector construct design, dose and animal age(*53*). The meta-analysis revealed that the most severe DRG and spinal cord findings usually occur with intra-CSF route of administration, with lack of a no-observed-adverse-effect-level (NOAEL) established at any dose level above the minimal efficacious dose (1E13 vg/animal) for most transgenes(*53*). The underlying cellular mechanisms that result in degeneration and axonopathy of the DRG sensory nerves following administration of AAV gene therapies have not been fully characterized, however, evidence supports the hypothesis that cellular stress due to high expression of transgene RNA and/or proteins is involved (*39, 40*). Additionally, the clinical translation of DRG toxicity seen in NHPs is not well understood, with more research needed to understand the mechanisms of toxicity as well as methods to evaluate and mitigate potential sensory neuron toxicity in clinical investigations. Demonstration of proof-of-concept in the mouse model of neuropathic pain as well as the 1-month NHP DRF study support initiation of an IND-enabling toxicology study to further characterize safety and select AAV9-hZFR dose levels for a potential Phase 1 clinical study in patients with peripheral neuropathy.

Peripheral neuropathies are considered chronic disorders as neuronal damage is generally not considered reversible because neurons lack regenerative capacity (*54*). As a result, genomic medicine approaches could potentially provide a long lasting and efficacious treatment for such chronic conditions. The efficacy of genomic medicine targeting *Scn9a* has been illustrated before in preclinical studies. For instance, targeting *Scn9a* RNA with shRNA or antisense oligonucleotides (ASOs) successfully reduced pain in pain rodent models (*18, 21*). However, targeting RNA is often associated with continuous treatment, off-target toxicity effects and insufficient target interactions (*55*), which makes it unsuitable for long-lasting effectiveness in the clinic for indications such as peripheral neuropathies. AAV-mediated delivery often results in a long-lasting effect in neurons (*56, 57*), which makes this delivery approach better suited for treatment of peripheral neuropathies. AAV9-delivered CRISPR-Cas9 has been used successfully in multiple mouse pain models (*20*), however, the large size makes it challenging for sufficient AAV9 packaging for large scale production. Unlike CRISPR-Cas9, ZFRs are small and compact, allowing for efficient AAV packaging. Additionally, as mentioned above, ZFRs are human-derived proteins and as such are not anticipated to provoke an immune response as anticipated with CRISPR-Cas9 bacterial components, which makes ZFRs a suitable approach for targeting the DRG neurons for treatment of peripheral neuropathies.

In conclusion, we have demonstrated that human or mice *SCN9A* targeting ZFR can selectively, effectively and robustly represses the expression of the *SCN9A* gene encoding the Nav1.7 sodium channel *in vitro* and in DRG neurons following a single IT-L administration in both mice and/or cynomolgus monkeys. Proof-of-concept was demonstrated in a mouse neuropathic pain model, providing insight into ZFR expression and Nav1.7 repression levels in DRG sensory neurons needed for efficacy. The pharmacologic activity and safety profile in mice and cynomolgus monkeys support moving the human/NHP *SCN9A*-targeted ZFR into IND-enabling toxicology studies.

## MATEIRALS AND METHODS

### Human and mouse ZFRs design

DNA Sequence encoding chimeric cys2-his2 zinc finger proteins (ZFPs) coupled with a recognition helix were designed based on a backbone derived from the human ZFP Zif268/EGR1. The recognition helices were selected from those previously validated for on-target specificity to nucleotide triplets. From this, arrays of 5 or 6-fingers (backbone and recognition helices) targeting unique sequences of a 15 to 18 nucleotides found near transcript start sites of the *SCN9A/Scn9a* in human (hg38) and mouse (mm10) reference genomes were selected. Sites designed against the human genome were further restricted to regions with homology to *SCN9A* in the *M.fascicularis* (macFas5) reference genome. The target sequences for the mouse and NHP/human ZFRs are as follow:

Mouse ZFR (mZFR): GTGCGAGTGTGCGCCAGT

NHP/human ZFR (hZFR): GGTGGCGACGCTGTAGCC

### Study design: Dose range-finding pharmacology and toxicology study in NHPs

Naïve, cynomolgus monkeys (sourced from Cambodia) were purchased by CRL. The study design consists of 1 vehicle control group (n=2, 1 male and 1 female) and 3 dose groups for hZFR, 1E12, 1E13, and 9E13 vg/animal, (n=3/group, 2 males and 1 female/group). AAV-hZFR doses were given as a single IT-L administration on Day 1 and animals were observed for 28 days post administration. On Day 29, animals were placed under deep and unrecoverable anesthesia followed by a cardiac perfusion with ice cold RNase-free phosphate buffered saline. The safety parameters evaluated included clinical observations, body weights, clinical pathology which included full panel of hematology (red blood cell count, hemoglobin concentration, hematocrit, mean corpuscular volume, red blood cell distribution width, mean corpuscular hemoglobin concentration, mean corpuscular hemoglobin, reticulocyte count [absolute], platelet count, white blood cell count, neutrophil count (absolute), lymphocyte count [absolute], monocyte count [absolute], eosinophil count [absolute], basophil count [absolute], large unstained cells [absolute]), chemistry (alanine aminotransferase, aspartate aminotransferase, alkaline phosphatase, gamma-glutamyl transferase, lactate dehydrogenase, total bilirubin, urea nitrogen, creatinine, calcium, phosphorus, low density lipid [LDL], and high density lipid [HDL]), and coagulation(activated partial thromboplastin time, prothrombin time, and fibrinogen), necropsy observations, and microscopic evaluation of the following tissues: Adrenal gland, brain (frontal cortex, cerebellum, brain stem), DRGs (sacral, lumbar, thoracic, cervical), epididymis, heart, kidney, large intestine, liver, lung, lymph node (mandibular), olfactory bulb, ovary, pancreas, sciatic nerve, skeletal muscle, small intestine, spinal cord, spleen, stomach, testes, thymus, uterus/cervices, and any macroscopic abnormalities. The following organs were weighed: brain, epididymis, heart, kidney (pair), liver, ovary (pair), spleen, testes (pair), and uterus/cervices. For All tissues were preserved in 10% neutral buffered formalin, embedded in paraffin and processed to blocks. For microscopic evaluations, tissues were cut in 5 micron thick sections, mounted on glass slides and stained with hematoxylin and eosin. All slides were evaluated by a CRL board certified veterinary pathologist.

For pharmacology assessment, DRGs were collected from lumbar (L5 and L1), thoracic (T7, T6, and T1), and cervical (C6 and C5) levels and placed in prelabelled tubes and flash frozen in liquid nitrogen. The tubes were kept in a −80°C freezer until analysis.

For single-cell analysis, DRGs were collected and preserved in 10% neutralized buffered formalin at room temperature for 24 hours, then transferred to 70% ethanol and processed to block within 7 days from transfer to ethanol. DRGs were embedded in paraffin and proceed to slides for analysis.

### ZFR and *SCN9A* mRNA analysis for dose range-finding pharmacology and toxicology study in NHP

Each DRG tissue was transferred to 2 mL Eppendorf tubes containing 0.9 mL TRI Reagent (ThermoFisher Scientific) and two 3.2 mm steel beads (BioSpec Products) on ice. The tissues were lysed using a Qiagen TissueLyser at 4°C using the following parameters: 15 cycles, 90s duration, 25.1 frequency. After centrifugation, 105 μL of 1-bromo-3-chloropropane was added to each sample at room temperature. The samples were vortexed for 10s, incubated for 5 min at room temperature, centrifuged at 12,000 ×g for 10 min at 4°C. Aqueous phases corresponding to the same original spinal cord level were combined. Four hundred microliters (400 μL) of the aqueous phase from each sample was transferred to a well of a 96 deep-well plate. Two hundred microliters (200 μL) of isopropyl alcohol were added to each sample well containing the aqueous phase of the tissue lysate. Samples were shaken for 1 min at 600 rpm at room temperature. Ten microliters of MagMax magnetic beads (ThermoFisher Scientific) were added to each sample well and mixed briefly. A KingFisher Flex Purification System (ThermoFisher Scientific) and MagMax Total RNA Isolation kit (ThermoFisher Scientific) were used to isolate RNA from tissue lysates following the manufacturer’s instructions. Approximately 100 μL of eluted RNA were separated from magnetic beads using a magnetic stand by incubating for 5 min at room temperature. Total RNA concentration and quality were evaluated using a Lunatic UV-vis absorbance spectrometer (Unchained Labs). Reverse transcription was prepared using the High-Capacity cDNA Reverse Transcription Kit (Applied Biosystems). Custom TaqMan primer:probe assays and TaqMan 2x Universal PCR Master Mix (Applied Biosystems) were used to perform qPCR using a QuantStudio 6 Flex Real Time PCR Machine (Applied Biosystems). A portion of each reverse transcription reaction was used to carry out two independent qPCR reactions – first for the absolute quantification of ZFR mRNA and second for multiplexed detection of *SCN9A*, *EIF4A2* and *ATP5B* mRNA. ZFR mRNA copy values were derived from the ZFR standard curve. The obtained ZFR copy values were then divided by the total RNA mass (ng) taken into the qPCR reaction to determine ZFR copies per ng of total RNA. *SCN9A* mRNA fold change was calculated using 2DeltaDeltaCt method formula where sample DeltaCt was calculated first as a difference between *SCN9A* cycle threshold (Ct) and geometric mean of two housekeeping genes Ct and then subtracted by the mean DeltaCt of the assay quality control samples. *SCN9A* mRNA fold change was then further normalized by dividing the sample fold change values by the mean fold change value from the untreated animals.

### Study design: Pain assessments in SNI mouse model

Naïve C57BL/6 female mice (8/sex/group), approximately 6-10 weeks old, were purchased 1 week prior to the initiation of each study. Animals were randomly assigned to either the control (vehicle or sham) or a ZFR treatment groups. ZFR were administered at a dose of 8E11 vg/mouse. The vehicle, method, volume, and route of administration, in-life assessments, necropsy, euthanasia, and tissue collection and storage procedures were as described above for the ZFR on target assessment study.

To generate the model, all animals on Day -7 were deeply anesthetized under 2.5% isoflurane in O_2_. The left hind leg was shaved, disinfected, and a small incision was made on the skin. The surgical site was opened with blunt dissection to visualize the three distal branches of sciatic nerve. Two of the three distal branches of sciatic nerve (tibial and peroneal nerve) were axotomized while sparing one (sural nerve). The surgical site was sutured, and animals were observed for proper wound healing.

Pain assessment (mechanical- and cold-induced) was performed prior to surgery, one week after the surgery and just prior to dose administration on Day 1, then on Days 3, 8, 15, 22, and 28. The pre-surgery measurement was conducted to establish a normal pain baseline for all animals prior to their assignment to the study. Any animal with abnormal pain sensitivity was not used. Assessment of pain (mechanical- and cold-induced) 7 days post-surgery (just prior to dose administration) was conducted to assure functionality of the model and to obtain a model baseline as animals that underwent the full surgery should have hypersensitivity to pain. The study included a sham operated group (skin opened and closed but no nerve was cut) to establish that the act of cutting open the skin does not impact pain sensitivity, a vehicle control group, a Gabapentin (GBP) group, and the test article groups. GBP (50 mg/kg; intraperitoneally) was administered 1 hour prior to each assessment. The dose of GBP was not an anesthetic dose in mice and was selected to reduce but not abolish pain in mice at 0.5-1 hour post-dose (*58*).

For the mechanical-induced pain assessment animals were placed individually into small cages with a mesh bottom. A monofilament (Von Frey fibers) was applied perpendicularly to the ventral surface of the left hind paw delivering a constant pre-determined force until it buckled at which time it was removed. A response was considered positive if the animal exhibited any nocifensive behaviors, including brisk paw withdrawal, licking, or shaking of the paw, either during application of the stimulus or immediately after the filament was removed. Testing began with the response to a filament estimated to be close to the 50% withdrawal threshold. If there was no response, the next filament with a higher force was tested; if there was a response, the next lower force filament was tested. Five continuous readings were taken and later assessed to determine withdrawal threshold in grams. For the cold-induced pain assessment animals were placed on a metal plate after it was cooled to the desired temperature and the time taken to evoke nociceptive behavior such as flinches, shaking, or licking in the affected paw or jumping was manually scored as paw withdrawal latency (sec). For SmartCage assessment, animals were placed in the SmartCage system individually and assessed for any movement abnormalities on Day 28. Animals were monitored for 22 hours, and data was analyzed at 1 hour blocks.

### Mouse and NHP DRGs single-cell analysis

For the mouse DRGs, single-cell analysis was performed at Evotech. DRG tissue sections on slides were hybridized with a combination of two RNAscope probes, panZFP probe (ACD Bio: 851651-C2) and MmScn9a probe (ACD Bio: 313341), Following hybridization, sections were stained with an antibody against peripherin (Abcam: ab246502at) at 1:1000 dilution. DAPI was used in all sections to mark cellular nuclei. The ISH/IHC multiplexing was performed according to the CRO protocol. Images were acquired with an Axio Scan.Z1 slide scanner (Carl Zeiss Microscopy GmbH), using a 20x Plan-apochromat objective (0.8 NA) and a Hamamatsu Camera. LED intensity, exposure time and emission filter settings were kept constant across groups for the acquisition. Images were acquired with a pixel resolution of 0.326 x 0.326 µm², inspected in ZEN (v.3.5, Carl Zeiss Microscopy GmbH) and analyzed using custom-written scripts in Acapella Studio 5.1 (PerkinElmer Inc.) before compiling in Spotfire (v.10.3.3 TIBCO).

For the NHP DRGs, the single-cell analysis was performed at ACD Bio. The NHP lumbar DRG was hybridized first with guinea pig anti-NeuN at 1:200 (Millipore Sigma catalog no. ABN90P) and mouse anti-CGRP (Abcam catalog no. ab81887). Next, A pooled RNAscope™ target probe consisting of Mfa-SCN9A (ACD catalog no. 591588) and pan-ZFP-KRAB-C2 (ACD catalog no. 851658-C2) was then hybridized for 2 hours at 42°C, followed by a series of amplification steps and rinse steps using RNAscope™ Multiplex amplification reagents per manufacturer’s instructions (ACD catalog no. 322800). Finally, DAPI was incubated for 10 minutes at room temperature for nuclear staining. Whole-tissue multiplex imaging was performed at 40X resolution using a 3DHISTECH PANNORAMIC SCAN II digital slide scanner, equipped with SpGr-B, SpOr-B, Cy5.5 and Cy7 filters for visualization of Vivid™ 520, Vivid™ 570, Alexa Fluor® 647 and Opal™ 780 fluorophores, respectively.

### Statistical analysis

Statistical analysis was performed using GraphPad Prism version 7 (GraphPad Software, San Diego, CA, USA). The statistical test used for each experiment is provided in the associated methods and/or figure legend. In general, one-way ANOVA with Dunnett’s for multiple comparison was used when comparing three or more groups. Significance is indicated by *. The following standard abbreviations are used to reference *P* values: ns, not significant; \**P* < 0.05; \*\**P* < 0.01; \*\*\**P* < 0.001; \*\*\*\**P* < 0.0001.

## Acknowledgments

We thank Y. Santiago for primary neuron culture and D. Chung for help with Affymetrix analysis. We thank C. Gasper for facilitating the animal studies, A. Ledeboer for reviewing animal protocols, K. Kennard for managing the animal studies and J. Perez for biorepository of animal samples. We thank S. Xie and N. Yan from AfaSci (Redwood City, CA, USA) for conducting the SNI mouse study, and individuals at Charles River Laboratory (Reno, NV, USA) for conducting the NHP study. We thank T. Fieblinger, G. Cisbani, and F. Peters at Evotec (Hamburg, Germany) for conducting single cell RNAscope analysis on mouse tissues and L. Chatelain and F. Li from ACD (Newark, CA, USA) for conducting single cell RNAscope analysis on NHP tissues. We thank A. Resch and J. Shoffner for helpful discussions and A. Resch for program oversight. We thank A. Young, G. Davis, Y. Lu, G. Brasnjo, E. McNeil, L. Wilkie, N. Dubois-Stringfellow, P. Ramsey and A. Resch for reviewing the manuscript.

## Competing interests

M. Samie and J. Eshleman are inventors on a pending United States patent application related to this work. M. Samie, T. Parman, M. Jalan, J. Lee, P. Dunn, J. Eshleman, Y. Pan, M. Falaleeva, S. Hinkley, T. Chen, S. Bhardwaj, A. Ward, M. Trias, A. Chikere, M. Som, S. Yadav, K. Meyer, B. Zeitler, and A. Pooler are current employees of Sangamo Therapeutics, Inc. D. Baldwin Vidales, J. Holter, B. Jones, and J. Fontenot were employed by Sangamo Therapeutics, Inc. when this work was conducted.

## List of Supplementary Materials

Materials and Methods Fig S1 to S3

## SUPPLEMENTAL MATERIALS

### Materials and Methods

#### Human SK-N-MC neuroepithelial cell line

Human SK-N-MC lines were purchased from ATCC (Catalog #HTB-10) and cultured according to the manufacturer protocol. Cells were maintained in tissue culture flasks until confluency with Eagle’s MEM cell media. For each passage, SK-N-MC cells were resuspended in culture media, and a small aliquot was mixed 1:1 with trypan blue solution 0.4% (w/v) in phosphate-buffered saline (PBS) and cells were counted on the TC20 Automated Cell Counter (Bio-Rad; Cat No. 145-0102). For nucleofection 100,000 cells per well were used for nucleofection.

#### Human iPSC-derived GABAergic neuron

Human iPSC-derived GABAergic neurons were purchased from Cellular Dynamics International and plated onto poly-L-ornithine- and laminin-coated 96-well plates. After thawing each vial of neurons, cells were resuspended in culture media, and a small aliquot was mixed 1:1 with trypan blue solution 0.4% (w/v) in phosphate-buffered saline (PBS) and cells were counted on the TC20 Automated Cell Counter (Bio-Rad; Cat No. 145-0102). Neurons were then seeded at a density of 40,000 cells per well of 96-well plate or 250,000 cells per well of a 24-well plate for qPCR or Affymetrix analysis, respectively.

#### Human iPSC-derived sensory neurons

Human iPSC-derived sensory neuron progenitor cells were purchased from Axol Biosciences (NHCDR000848). Cells were cultured and differentiated on PDL coated plates according to the manufacturer protocol. Cells were differentiated for 10 days and then transduced with different AAV-ZFR construct at the indicated MOIs. Cells were harvested on day 18 and RNA was isolated, and RT-qPCR was performed for gene expression analysis.

For patient derived cells, the RCi001-A iPSC line was developed from a male individual with IEM (*59*). The cell line was purchased from the European Bank for induced pluripotent Stem Cells (EBiSC). For the control lines, a healthy iPSC line (NHCDR250422) was purchased from NINDS. Both lines differentiated to sensory neurons by Axol Biosciences by using Sendai reprogramming method. Cells were cultured and used similar to healthy progenitor cells per manufacturer protocol.

#### Mouse N2a neuroblastoma cell line

Mouse N2a neuroblastoma cells were purchased from ATCC (Catalog # CCL-131) and cultured according to the manufacturer protocol. Cells were maintained in tissue culture flasks until confluency with Eagle’s MEM cell media. For each passage, N2a cells were resuspended in culture media, and a small aliquot was mixed 1:1 with trypan blue solution 0.4% (w/v) in phosphate-buffered saline (PBS) and cells were counted on the TC20 Automated Cell Counter (Bio-Rad; Cat No. 145-0102). For nucleofection, 100,000 cells per well were used.

#### Mouse primary cortical neurons

Primary mouse cortical neurons (MCNs) were purchased from Gibco. Cells were plated onto poly-D-lysine-coated 24-well plates at 200,000 cells/well and maintained according to the manufacturer’s specifications using Gibco Neurobasal Medium containing GlutaMAX™ I supplement, B27 supplement, and penicillin/streptomycin. 48 hours after plating (at DIV2), cells were transduced with AAV6-ZFRs (at the 1.00E4 MOIs and harvested 7 days later (at DIV9; 50% media exchanges performed every 2-3 days) followed by RNA isolation and Affymetrix analysis.

#### ZFR mRNA nucleofection

Cells were plated on 96-well plates and were resuspended in Amaxa^®^ SF solution. The cells were then mixed with ZFR mRNA at 3 doses: 100, 300, 1000 ng, and transferred to Amaxa^®^ shuttle plate wells. The cells were transfected using the Amaxa^®^ Nucleofector^®^ device (Lonza; program CM-137). Eagle’s MEM cell media was added to each well of the plate. The cells were transferred to a 96-well tissue culture plate and incubated at 37°C for 20 hours. Cells were harvested after 20 hours and processed for RT-qPCR analysis.

#### AAV packaging and production of ZFRs

Recombinant adeno-associated viral vectors (rAAV) were generated by the triple transfection method. Briefly, HEK293 cells were plated in ten-layer CellSTACK chambers (Corning, Acton, MA) and grown for three days to a density of 80%. Three plasmids – (i) an AAV Helper plasmid containing the Rep and Cap genes, (ii) an Adenovirus Helper plasmid containing the adenovirus helper genes, and (iii) a transgene plasmid containing the sequence to be packaged flanked by AAV2 inverted terminal repeats were transfected into the cells using calcium phosphate. After three days, the cells were harvested. The cells were then lysed by three rounds of freeze/thaw and the cell debris was removed by centrifugation. The rAAV was precipitated using polyethylene glycol. After resuspension, the virus was purified by ultracentrifugation overnight on a cesium chloride gradient. The virus was formulated by dialysis and then filter-sterilized. After adjusting the titer (vector genomes/mL; vg/mL) of all AAV batches by dilution with PBS + 0.001% Pluronic F-68, the AAVs were aliquoted to single use doses and stored at −80°C until use. No samples were frozen after they were thawed.

#### *In vitro* transduction of iPSC-derived neurons with AAV6-ZFRs

The cells were transfected with AAV6-ZFRs at different MOIs 48 hours after plating and maintained for up to 4 weeks (50-75% media changes performed every 3-5 days). For RT-qPCR analysis, Human iPSC-derived GABAergic and sensory neurons were transduced with the following MOIs: 3.00E3, 1.00E4, 3.00E4,1.00E5, and 3.00E5 vg/cell. Cells were harvested at the end of the experimental period, RNA was isolated, and RT-qPCR was performed. For Affymetrix analysis, the cells were transfected with 1.00E5 vg/cell 48 hours after plating and harvested 19 days after viral transfection.

#### RNA isolation and RT-qPCR from cells

After each time point for each cell line, cells were harvested and lysed and reverse transcription was performed using a Cells2CT Custom kit (Cat# 4402955C001) following the manufacturer’s instructions. TaqMan quantitative polymerase chain reaction (qPCR) was used to measure the expression levels of different genes, which were normalized to the geometric mean of the expression levels of the housekeeping genes *Atp5b* and *Eif4a2*. RT-qPCR was performed using Bio-Rad CFX384 thermal cyclers. cDNA was diluted 10-fold in nuclease-free water, and 4 μL of diluted cDNA were added to each 10 μL PCR reaction. Plates were prepared using Tecan automated liquid handling robotics. Each sample was assayed in technical quadruplicate. The following primer/probes from IDT were used for RT-qPCR analysis for different genes:

Primers targeting mouse genes:

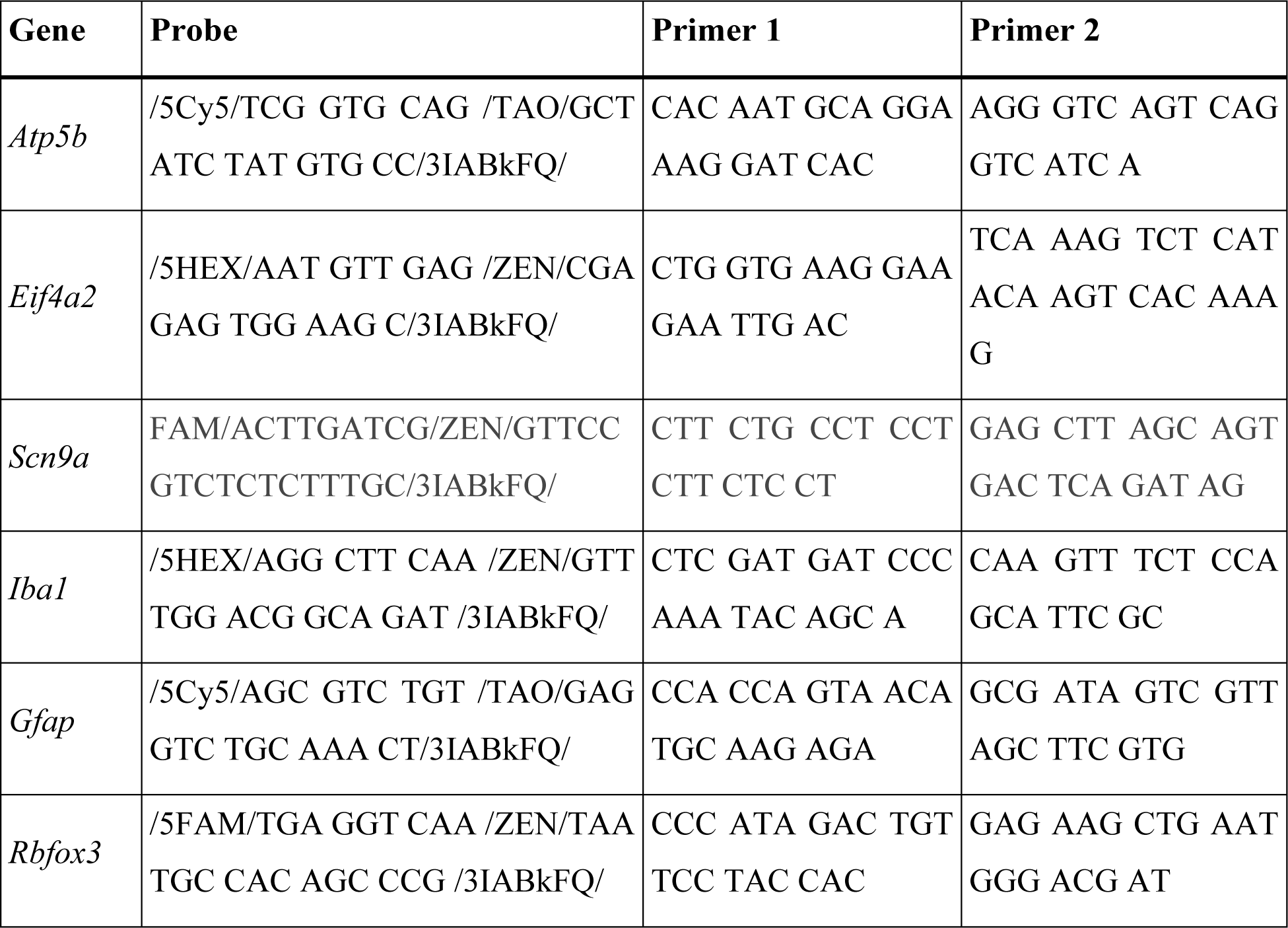

Primers targeting human genes:

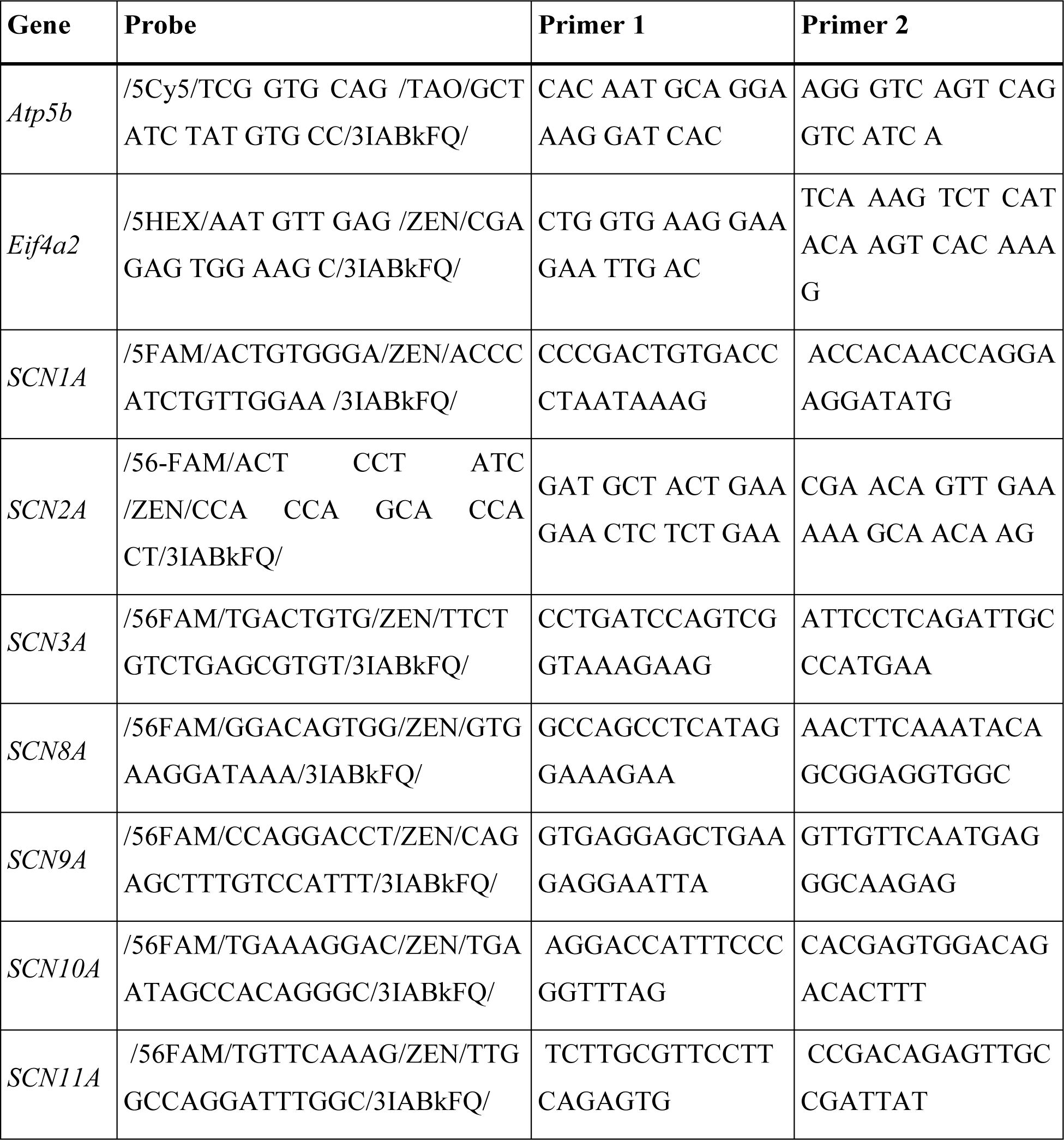

Primers targeting NHP genes:

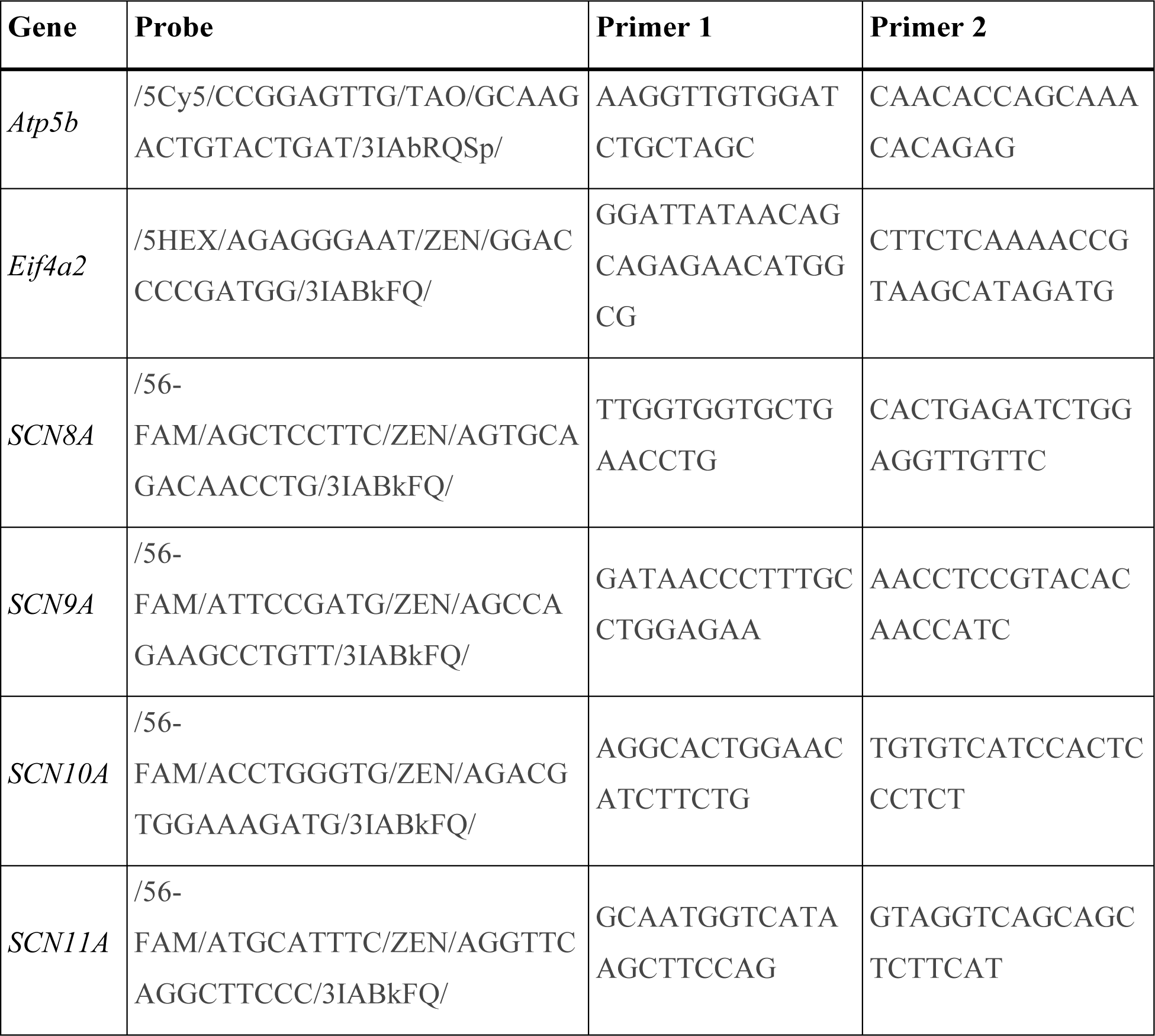

ZFR expression

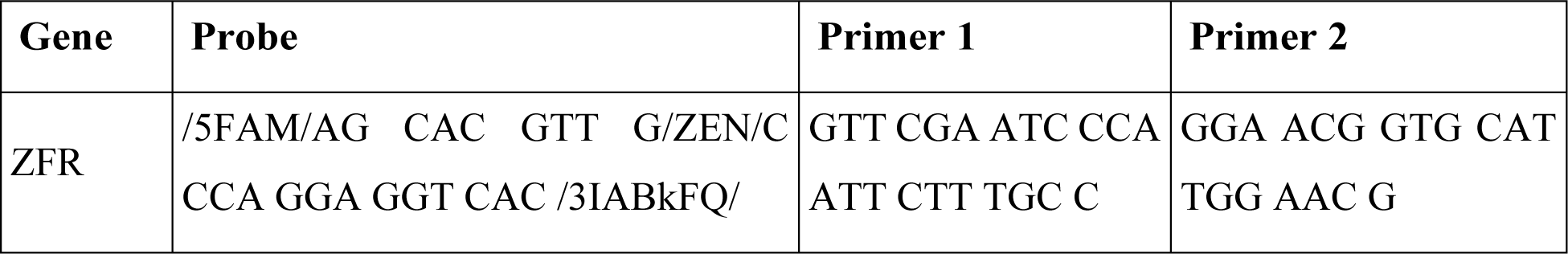

#### Affymetrix analysis for the evaluation of off-targets

To evaluate the potential off-target impact of the AAV6-ZFRs on global gene expression, we performed microarray experiments (Clariom™ S Assay HT, mouse (Thermofisher Cat. #902972) for mouse primary cortical neurons) on total RNA isolated from primary mouse cortical neurons transduced with AAV6-ZFRs. For each replicate (50 ng of total RNA) was processed according to the manufacturer’s protocol for sample preparation, hybridization, fluidics, and scanning (Affymetrix MTA1.0 GeneChip arrays, Affymetrix). Fold change analysis was performed using Transcriptome Analysis Console 4.0 (Affymetrix) software. AAV6-ZFR-treated samples were compared to samples treated with a non–Scn9a-targeted AAV6-ZFR. Change in gene expression is reported for transcripts (probe sets) with a >2-fold difference in mean signal relative to control and with the FDR-corrected p-values set to <0.05.

#### Ethical statement for animal studies

Mouse studies were conducted at Afasci (Redwood City, CA, USA) under a protocol approved by the Institutional Animal Care and Use Committee. The NHP study was conducted at Charles River Laboratories, Inc. (CRL, Reno, NV, USA) which is an Association for Assessment and Accreditation of Laboratory Animal Care International (AAALAC) accredited facility under an Institutional Animal Care and Use Committee approved protocol. All studies were performed in conformance with the U.S. Public Health Policy on the Care and Use of Animals as defined in the Guide to the Care and Use of Animals. Species specific standard procedures and conditions for animal care, housing, access to water and food, environment, and room maintenance were used. All other procedures were performed in accordance with laboratory standard operating procedures and/or established laboratory best practices.

#### mZFR on target assessment study in mice

Naïve C57BL/6 female mice (8/sex/group), approximately 5-7 weeks old were purchased from Charles River Laboratories Inc. (CRL), one week prior to the initiation of each study. Animals were randomly assigned to either control or test article treatment groups. ZFR was administered as a single bolus dose of 2E11 vg/ animal via IT-L into the L4-L5 or L5-L6 intervertebral space on Day 1 and necropsy occurred on Day 22. ZFR was formulated in an AAV formulation buffer-P [1X Dulbecco’s Phosphate-buffered saline (DPBS) (+Ca/+Mg) + 0.001% Pluronic F-68, pH 7.2]. Volume of administration was 10 μL. Mice were excluded from the study and replaced if no tail flick was observed to indicate successful IT-L administration. Animals were observed for mortality/morbidity and general health at cageside daily and detailed clinical observation once weekly. Body weights were measured predose and once weekly. Mice were euthanized by isoflurane/oxygen anesthesia followed by whole-body transcranial perfusion with RNase-free 0.9% saline. Gross necropsy observations were performed. Three pairs of DRGs were collected from each of the cervical, thoracic, and lumbar regions and transferred to three separate 1.5 mL RNase free Eppendorf tubes. RNALater was added and tubes were refrigerated (1 to 8°C) for 24 hours. Following 24-hour incubation, tissues were removed from RNALater and transferred to labeled RNase free Eppendorf tubes. The tubes containing the tissue were then frozen on dry ice and kept in a −80°C freezer until analysis.

#### Mouse tissue processing and RNA isolation from the mouse DRGs

A total of 0.6 mL TRI reagent (Thermo Fisher) and two 3.2 mm steel beads (BioSpec Products) were added to each 1.5 mL Eppendorf tube on ice. The tissue was lysed using a Qiagen Tissue-Lyser (catalog number MW 3000) at 4°C using the following parameters: 5 cycles, 90 s duration, 25.1 frequency. After brief centrifugation, 70 μL of 1-bromo-3-chloropropane (BCP) was added to each sample at room temperature. The samples were vortexed for 10 s, centrifuged at 12,000 ×g for 10 min at 4°C, and 120 μL of the aqueous phase from each sample was transferred to wells of a 96-well plate. Sixty microliters of isopropyl alcohol and 10 μL of MagMax magnetic beads (Thermo Scientific) were added to each sample well containing the aqueous phase samples. A Kingfisher 96 robot (Thermo Scientific) and the MagMax kit (Thermo Fisher) were used to isolate RNA from the tissue lysate following the manufacturer’s instructions. One hundred microliters of the eluted RNA were separated from the magnetic beads using a magnetic stand. RNA yield and quality were evaluated using a Nanodrop 8000 instrument (Thermo Scientific).

**Figure S1.**
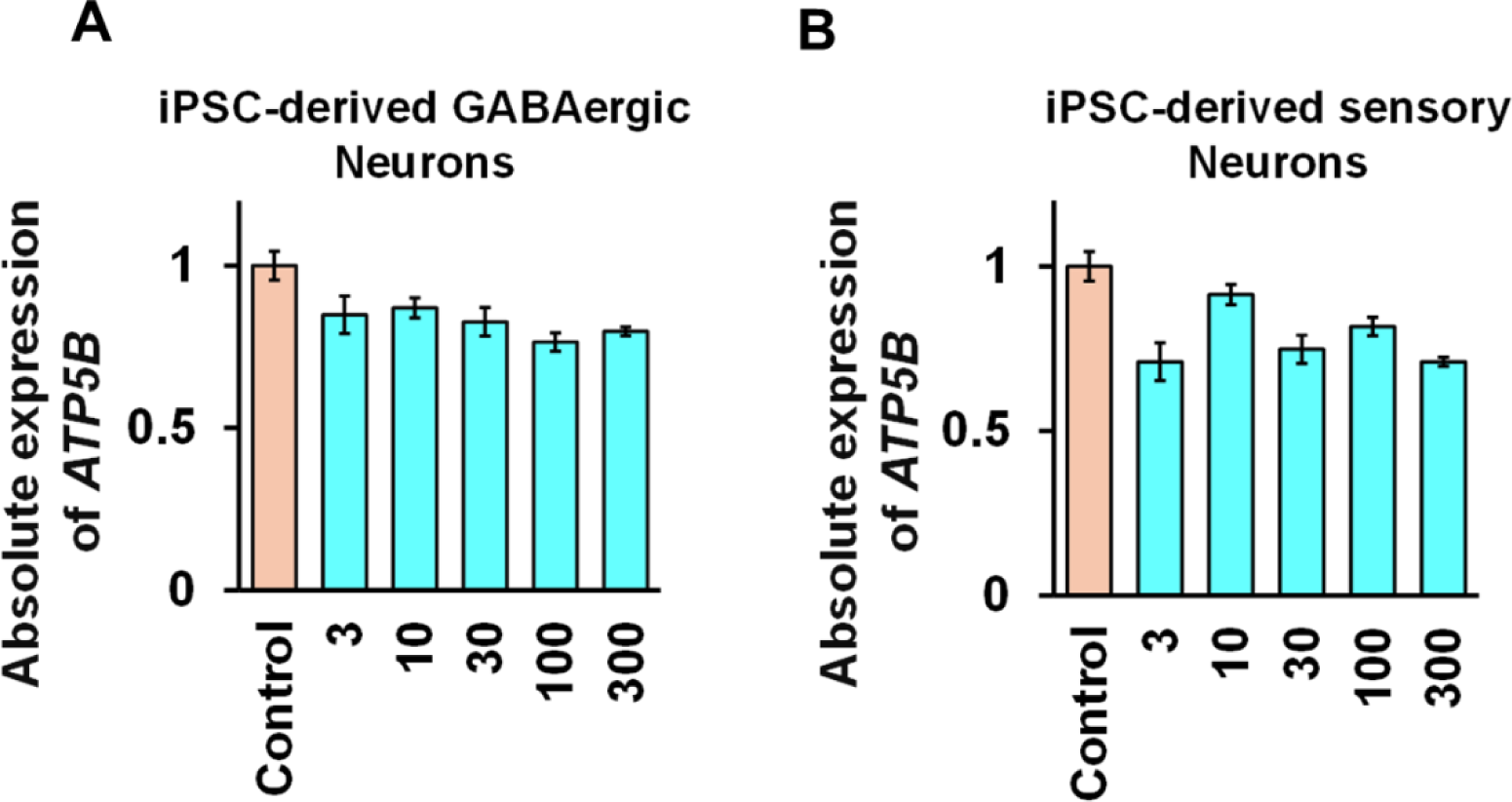
hZFR potent repression of *SCN9A* is not associated with decrease in housekeeping gene expression. a. mRNA expression of *ATG5B* was evaluated following hZFR treatment at different doses in iPSC-derived GABAergic neurons, Mean ± SD. b. mRNA expression of *ATG5B* was evaluated following hZFR treatment at different doses in iPSC-derived sensory neurons, Mean ± SD.

**Figure S2.**
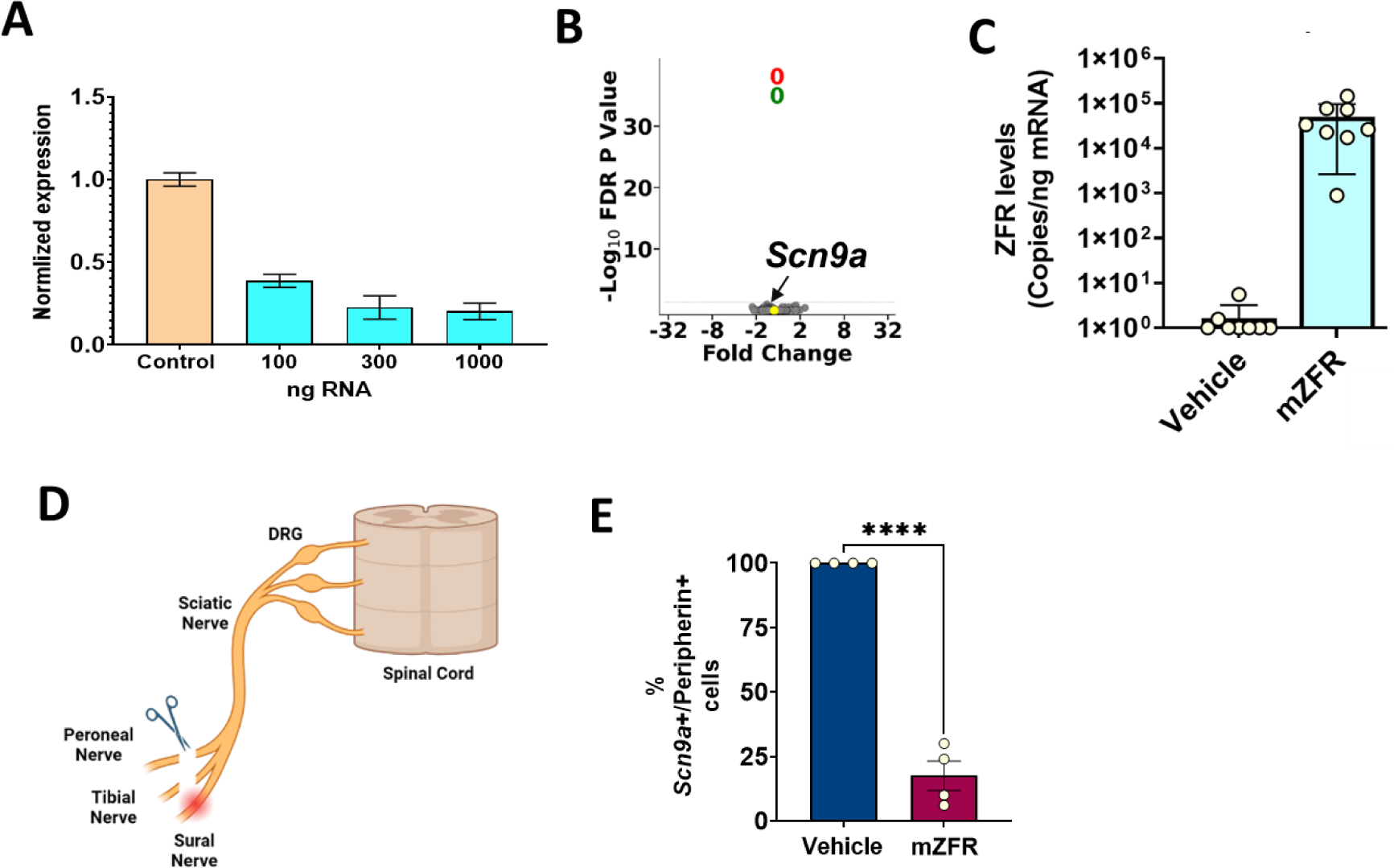
mZFR potently and specifically represses *Scn9a* in mouse neurons. **(A)** Significant reduction of *Scn9a* transcriptional levels in a dose-dependent manner in MCNs following AAV6-mZFP transduction. Doses (MO1) 1E3, 3E3, 1E4, 3E4, and 1E5. Mean ± SD. **(B)** mRNA microarray assessment (volcano plots) of MCNs 7 days post-transduction with mZFR packed into AAV6 and delivered to cells at 3E4 MOI. Each dot represents the mean fold change compared to control-treated cells for a given gene (x value) and the associated P value (y value). Gene expression profiles are calculated based on FDR adjusted P value >0.05. 6 biological replicates. **(C)** mZFR mRNA expression in lumbar DRG in mice 4 weeks post IT-L administration compared to the vehicle group. Each circle represents an animal. **(D)** Schematic diagram illustrating the spared nerve injury (SNI) model. SNI model is made surgically by damaging two of the three terminal branches of the sciatic nerve (tibial and common peroneal nerves) leaving the sural nerve intact. This leads to persistent and reliable tactile hypersensitivity under the skin area of the sural nerve. **(E)** Reduction of *Scn9a* mRNA in the peripherin-positive cells. Each circle represents a single DRG, and absolute values (as median *Scn9a* spot density per cell) are shown. Mean ± SEM, ordinary one-way ANOVA, ** P < 0.01 *** P < 0.001 **** P < 0.0001 Compared with Vehicle.

**Figure S3.**
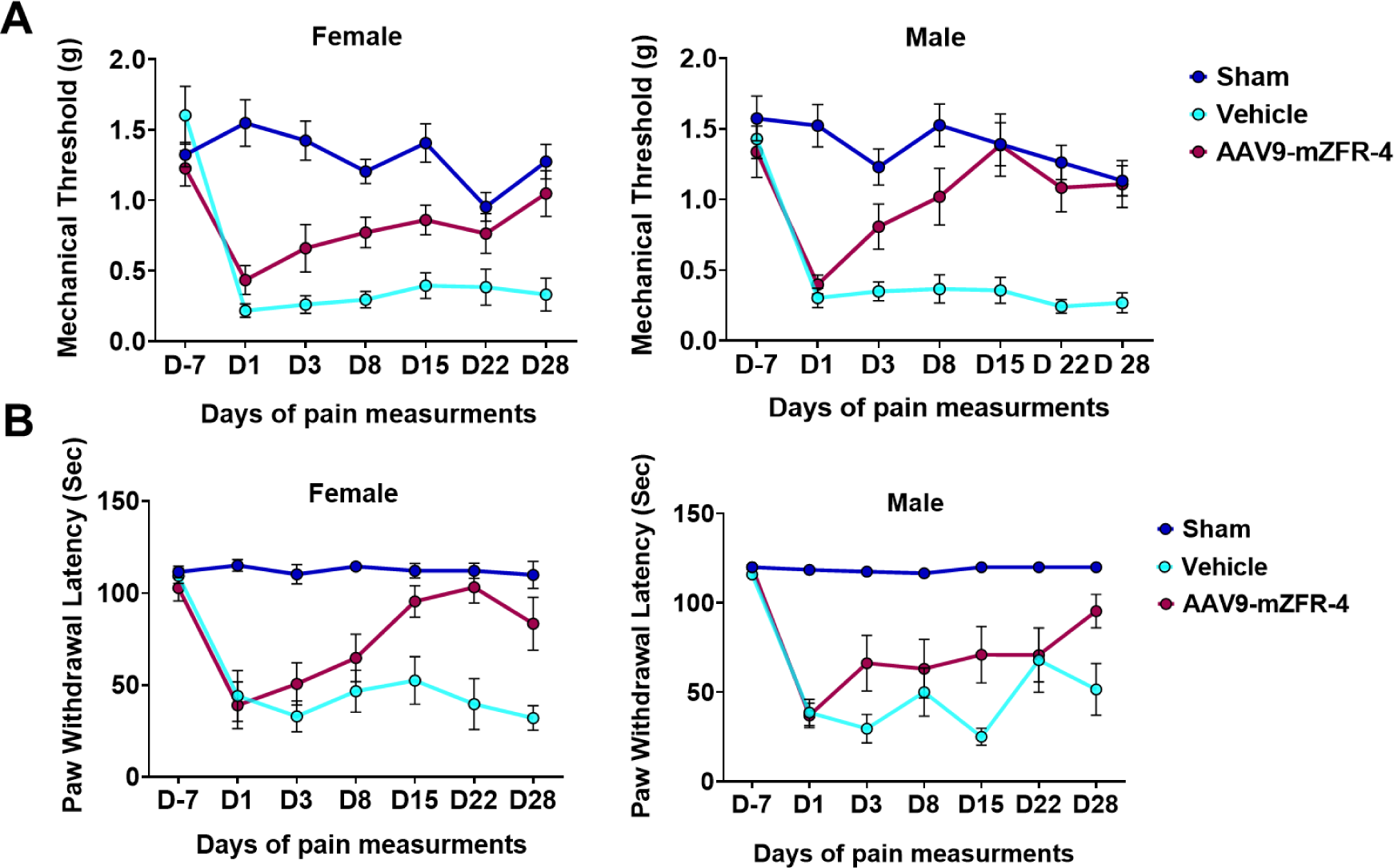
AAV9-mZFR is able to rescue the pain phenotype 1 week after IT-L injection in SNI neuropathic pain model. Longitudinal assessment of mechanical (A) and cold (B) induced pain responses in female and male animals following AAV9-mZFR treatment at Day -7 (D-7), 1, 3, 8, 15, 22, and 28. Both mechanical and cold induced pain thresholds are increased as early as Day 3 post IT-L injection and reach the same level of sham treated animals at Day 28

## References and Notes

1. N. Attal, M. Lanteri-Minet, B. Laurent, J. Fermanian, D. Bouhassira, The specific disease burden of neuropathic pain: results of a French nationwide survey. Pain 152, 2836–2843 (2011).

2. N. Torrance, B. H. Smith, M. I. Bennett, A. J. Lee, The epidemiology of chronic pain of predominantly neuropathic origin. Results from a general population survey. J Pain 7, 281–289 (2006).

3. T. S. Jensen et al., A new definition of neuropathic pain. Pain 152, 2204–2205 (2011).

4. A. B. O’Connor, Neuropathic pain: quality-of-life impact, costs and cost effectiveness of therapy. Pharmacoeconomics 27, 95–112 (2009).

5. B. Bingham, S. K. Ajit, D. R. Blake, T. A. Samad, The molecular basis of pain and its clinical implications in rheumatology. Nat Clin Pract Rheumatol 5, 28–37 (2009).

6. R. S. Y. Ma et al., Voltage gated sodium channels as therapeutic targets for chronic pain. J Pain Res 12, 2709–2722 (2019).

7. M. de Lera Ruiz, R. L. Kraus, Voltage-Gated Sodium Channels: Structure, Function, Pharmacology, and Clinical Indications. J Med Chem 58, 7093–7118 (2015).

8. F. H. Yu, W. A. Catterall, Overview of the voltage-gated sodium channel family. Genome Biol 4, 207 (2003).

9. J. P. Drenth, S. G. Waxman, Mutations in sodium-channel gene SCN9A cause a spectrum of human genetic pain disorders. J Clin Invest 117, 3603–3609 (2007).

10. T. R. Cummins, S. D. Dib-Hajj, S. G. Waxman, Electrophysiological properties of mutant Nav1.7 sodium channels in a painful inherited neuropathy. J Neurosci 24, 8232–8236 (2004).

11. Y. Yang et al., Mutations in SCN9A, encoding a sodium channel alpha subunit, in patients with primary erythermalgia. J Med Genet 41, 171–174 (2004).

12. C. R. Fertleman et al., SCN9A mutations in paroxysmal extreme pain disorder: allelic variants underlie distinct channel defects and phenotypes. Neuron 52, 767–774 (2006).

13. J. J. Cox et al., An SCN9A channelopathy causes congenital inability to experience pain. Nature 444, 894–898 (2006).

14. S. D. Dib-Hajj, T. R. Cummins, J. A. Black, S. G. Waxman, From genes to pain: Na v 1.7 and human pain disorders. Trends Neurosci 30, 555–563 (2007).

15. S. D. Dib-Hajj, Y. Yang, J. A. Black, S. G. Waxman, The Na(V)1.7 sodium channel: from molecule to man. Nat Rev Neurosci 14, 49–62 (2013).

16. L. A. McDermott et al., Defining the Functional Role of Na(V)1.7 in Human Nociception. Neuron 101, 905–919 e908 (2019).

17. S. Ahmad et al., A stop codon mutation in SCN9A causes lack of pain sensation. Hum Mol Genet 16, 2114–2121 (2007).

18. W. Cai et al., shRNA mediated knockdown of Nav1.7 in rat dorsal root ganglion attenuates pain following burn injury. BMC Anesthesiol 16, 59 (2016).

19. A. Mohan et al., Antisense oligonucleotides selectively suppress target RNA in nociceptive neurons of the pain system and can ameliorate mechanical pain. Pain 159, 139–149 (2018).

20. A. M. Moreno et al., Long-lasting analgesia via targeted in situ repression of Na(V)1.7 in mice. Sci Transl Med 13, (2021).

21. J. Pan, X. J. Lin, Z. H. Ling, Y. Z. Cai, Effect of down-regulation of voltage-gated sodium channel Nav1.7 on activation of astrocytes and microglia in DRG in rats with cancer pain. Asian Pac J Trop Med 8, 405–411 (2015).

22. C. G. Faber et al., Efficacy and safety of vixotrigine in idiopathic or diabetes-associated painful small fibre neuropathy (CONVEY): a phase 2 placebo-controlled enriched-enrolment randomised withdrawal study. EClinicalMedicine 59, 101971 (2023).

23. A. Dormer et al., A Review of the Therapeutic Targeting of SCN9A and Nav1.7 for Pain Relief in Current Human Clinical Trials. J Pain Res 16, 1487–1498 (2023).

24. D. A. Eagles, C. Y. Chow, G. F. King, Fifteen years of Na(V) 1.7 channels as an analgesic target: Why has excellent in vitro pharmacology not translated into in vivo analgesic efficacy? Br J Pharmacol 179, 3592–3611 (2022).

25. K. Kingwell, Nav1.7 withholds its pain potential. Nat Rev Drug Discov, (2019).

26. J. V. Mulcahy et al., Challenges and Opportunities for Therapeutics Targeting the Voltage-Gated Sodium Channel Isoform Na(V)1.7. J Med Chem 62, 8695–8710 (2019).

27. M. Imbeault, P. Y. Helleboid, D. Trono, KRAB zinc-finger proteins contribute to the evolution of gene regulatory networks. Nature 543, 550–554 (2017).

28. B. Zeitler et al., Allele-selective transcriptional repression of mutant HTT for the treatment of Huntington’s disease. Nat Med 25, 1131–1142 (2019).

29. S. Wegmann et al., Persistent repression of tau in the brain using engineered zinc finger protein transcription factors. Sci Adv 7, (2021).

30. Y. E. Tak et al., Genome-wide functional perturbation of human microsatellite repeats using engineered zinc finger transcription factors. Cell Genom 2, (2022).

31. J. F. Margolin et al., Kruppel-associated boxes are potent transcriptional repression domains. Proc Natl Acad Sci U S A 91, 4509–4513 (1994).

32. B. L. Ellis et al., A survey of ex vivo/in vitro transduction efficiency of mammalian primary cells and cell lines with Nine natural adeno-associated virus (AAV1-9) and one engineered adeno-associated virus serotype. Virol J 10, 74 (2013).

33. M. Richner, O. J. Bjerrum, A. Nykjaer, C. B. Vaegter, The spared nerve injury (SNI) model of induced mechanical allodynia in mice. J Vis Exp, (2011).

34. S. Shiers, R. M. Klein, T. J. Price, Quantitative differences in neuronal subpopulations between mouse and human dorsal root ganglia demonstrated with RNAscope in situ hybridization. Pain 161, 2410–2424 (2020).

35. C. Hinderer et al., Severe Toxicity in Nonhuman Primates and Piglets Following High-Dose Intravenous Administration of an Adeno-Associated Virus Vector Expressing Human SMN. Hum Gene Ther 29, 285–298 (2018).

36. J. Hordeaux et al., Toxicology Study of Intra-Cisterna Magna Adeno-Associated Virus 9 Expressing Iduronate-2-Sulfatase in Rhesus Macaques. Mol Ther Methods Clin Dev 10, 68–78 (2018).

37. J. Hordeaux et al., Toxicology Study of Intra-Cisterna Magna Adeno-Associated Virus 9 Expressing Human Alpha-L-Iduronidase in Rhesus Macaques. Mol Ther Methods Clin Dev 10, 79–88 (2018).

38. J. B. Rosenberg et al., Safety of Direct Intraparenchymal AAVrh.10-Mediated Central Nervous System Gene Therapy for Metachromatic Leukodystrophy. Hum Gene Ther 32, 563–580 (2021).

39. N. Buss et al., Characterization of AAV-mediated dorsal root ganglionopathy. Mol Ther Methods Clin Dev 24, 342–354 (2022).

40. J. Hordeaux et al., MicroRNA-mediated inhibition of transgene expression reduces dorsal root ganglion toxicity by AAV vectors in primates. Sci Transl Med 12, (2020).

41. A. H. Doth, P. T. Hansson, M. P. Jensen, R. S. Taylor, The burden of neuropathic pain: a systematic review and meta-analysis of health utilities. Pain 149, 338–344 (2010).

42. L. Colloca et al., Neuropathic pain. Nat Rev Dis Primers 3, 17002 (2017).

43. N. Attal, Pharmacological treatments of neuropathic pain: The latest recommendations. Rev Neurol (Paris) 175, 46–50 (2019).

44. N. B. Finnerup et al., Pharmacotherapy for neuropathic pain in adults: a systematic review and meta-analysis. Lancet Neurol 14, 162–173 (2015).

45. N. Nishikawa, M. Nomoto, Management of neuropathic pain. J Gen Fam Med 18, 56–60 (2017).

46. E. D. McNicol, A. Midbari, E. Eisenberg, Opioids for neuropathic pain. Cochrane Database Syst Rev 2013, CD006146 (2013).

47. S. G. Waxman, S. D. Dib-Hajj, Na(V)1.7: A central role in pain. Neuron 111, 2615–2617 (2023).

48. A. McDonnell et al., Efficacy of the Nav1.7 blocker PF-05089771 in a randomised, placebo-controlled, double-blind clinical study in subjects with painful diabetic peripheral neuropathy. Pain 159, 1465–1476 (2018).

49. G. Goodwin, S. B. McMahon, The physiological function of different voltage-gated sodium channels in pain. Nat Rev Neurosci 22, 263–274 (2021).

50. J. C. Chambers et al., Genetic variation in SCN10A influences cardiac conduction. Nat Genet 42, 149–152 (2010).

51. R. Coppini, C. Ferrantini, NaV1.8: a novel contributor to cardiac arrhythmogenesis in heart failure. Cardiovasc Res 114, 1691–1693 (2018).

52. A. O. Verkerk et al., Functional Nav1.8 channels in intracardiac neurons: the link between SCN10A and cardiac electrophysiology. Circ Res 111, 333–343 (2012).

53. J. Hordeaux et al., Adeno-Associated Virus-Induced Dorsal Root Ganglion Pathology. Hum Gene Ther 31, 808–818 (2020).

54. S. Blackshaw, Why Has the Ability to Regenerate Following CNS Injury Been Repeatedly Lost Over the Course of Evolution? Front Neurosci 16, 831062 (2022).

55. C. Rinaldi, M. J. A. Wood, Antisense oligonucleotides: the next frontier for treatment of neurological disorders. Nat Rev Neurol 14, 9–21 (2018).

56. Y. Chu, R. T. Bartus, F. P. Manfredsson, C. W. Olanow, J. H. Kordower, Long-term post-mortem studies following neurturin gene therapy in patients with advanced Parkinson’s disease. Brain 143, 960–975 (2020).

57. P. Hadaczek et al., Eight years of clinical improvement in MPTP-lesioned primates after gene therapy with AAV2-hAADC. Mol Ther 18, 1458–1461 (2010).

58. N. Kusunose et al., Molecular basis for the dosing time-dependency of anti-allodynic effects of gabapentin in a mouse model of neuropathic pain. Mol Pain 6, 83 (2010).

59. L. Cao et al., Pharmacological reversal of a pain phenotype in iPSC-derived sensory neurons and patients with inherited erythromelalgia. Sci Transl Med 8, 335ra356 (2016).

